# Distinctive and complementary roles of default mode network subsystems in semantic cognition

**DOI:** 10.1101/2023.10.03.560166

**Authors:** Ximing Shao, Katya Krieger-Redwood, Meichao Zhang, Paul Hoffman, Lucilla Lanzoni, Robert Leech, Jonathan Smallwood, Elizabeth Jefferies

## Abstract

The default mode network (DMN) typically deactivates to external tasks, yet supports semantic cognition. It comprises medial temporal (MT), core, and fronto-temporal (FT) subsystems, but its functional organisation is unclear: the requirement for perceptual coupling versus decoupling, input modality (visual/verbal), type of information (social/spatial) and control demands all potentially affect its recruitment. We examined the effect of these factors on activation and deactivation of DMN subsystems during semantic cognition, across four task-based functional magnetic resonance imaging (fMRI) datasets, and localised these responses in whole-brain state space defined by gradients of intrinsic connectivity. FT showed activation consistent with a central role across domains, tasks and modalities, although it was most responsive to abstract, verbal tasks; this subsystem uniquely showed more ‘tuned’ states characterised by increases in both activation and deactivation when semantic retrieval demands were higher. MT also activated to both perceptually-coupled (scenes) and decoupled (autobiographical memory) tasks, and showed stronger responses to picture associations, consistent with a role in scene construction. Core DMN consistently showed deactivation, especially to externally-oriented tasks. These diverse contributions of DMN subsystems to semantic cognition were related to their location on intrinsic connectivity gradients: activation was closer to sensory-motor cortex than deactivation, particularly for FT and MT, while activation for core DMN was distant from both visual cortex and cognitive control. These results reveal distinctive yet complementary DMN responses: MT and FT support different memory-based representations that are accessed externally and internally, while deactivation in core DMN is associated with demanding, external semantic tasks.

**Significance Statement:** We delineate the functional organisation of DMN in semantic cognition, examining effects of perceptual coupling versus decoupling, input modality (visual/verbal), domain (social/spatial) and control demands across DMN subsystems in four fMRI datasets. These subsystems played complementary roles in semantic cognition related to their locations on gradients of intrinsic connectivity. Medial temporal and frontotemporal subsystems supported visuospatial and abstract conceptual information respectively, across both internally and externally-focussed tasks, while deactivation in core DMN was associated with focussed and externally-oriented semantic states. We conclude that both content and process are relevant to the functional architecture of DMN in semantic cognition.

## Introduction

Default mode network (DMN) – with distributed components across medial and lateral frontal, parietal and temporal cortex – was initially characterized as task negative: it deactivates in attention-demanding tasks (Raichle et al., 2001; Greicius et al., 2003) and shows anti-correlated intrinsic connectivity with dorsal attention network (Greicius et al., 2003; Fox et al., 2005). However, DMN can couple with visual inputs (Zhang et al., 2019; Zhang et al., 2022) and it supports both perceptually-decoupled states (Christoff et al., 2009; Konu et al., 2020) and aspects of externally-oriented cognition, including semantic processing (Binder et al., 2009; Krieger-Redwood et al., 2016) and working memory (Spreng et al., 2014; Vatansever et al., 2015; Murphy et al., 2018).

A contemporary topographical perspective suggests these diverse functions might reflect DMN’s maximal distance from unimodal cortex (Margulies et al., 2016; Smallwood et al., 2021). This view emerges from whole-brain decompositions of intrinsic connectivity, termed ‘gradients’, which capture key features of cortical organization (Margulies et al., 2016; Hong et al., 2020; McKeown et al., 2020). The principal gradient, explaining the most variance, captures the separation of heteromodal DMN from sensory-motor cortex (Margulies et al., 2016) and correlates with physical distance from sensory-motor landmarks on the cortical surface. The second gradient relates to the distinction between visual and motor cortex, while the third gradient reflects the division within heteromodal cortex between controlled and less controlled responses. DMN’s position on the principal gradient far from sensory-motor cortex might support perceptually decoupled states. However, DMN’s position at the end of multiple processing streams might also facilitate the coordination and abstraction of higher-order representations.

DMN also contains subsystems associated with different cognitive processes (Andrews-Hanna, 2012; Andrews-Hanna et al., 2014): (i) Medial temporal (MT) DMN is linked to episodic memory and scene construction; (ii) core DMN with self-referential and perceptually-decoupled cognition; while (iii) fronto-temporal (FT) DMN is thought to support abstract, semantic and social cognition (Andrews-Hanna, 2012; Andrews-Hanna and Grilli, 2021). These subnetworks suggest a complex functional organisation, the principles of which are still not fully understood. While tasks eliciting FT activation are often semantic and tasks eliciting MT and core activation often involve episodic retrieval, there are multiple confounds in this contrast: semantic cognition is typically more perceptually coupled (involving access to meaning from visual inputs), abstract (involving verbal or categorical representations, as opposed to reconstructions of places and events) and controlled (involving more ambiguous decisions about information in the absence of recent exposure, cf. Vatansever et al., 2021). These dimensions might affect activation and deactivation in DMN subsystems in distinct ways, with all three subnetworks supporting semantic cognition if task demands are configured appropriately (Humphreys et al., 2015; Krieger-Redwood et al., 2016; Vatansever et al., 2021). Functional differences between the DMN subnetworks might also reflect the location of activating and deactivating voxels in gradient space, as the gradients relate to these frequently confounded cognitive dimensions, including abstraction (principal gradient), the balance of sensory-motor inputs (second gradient) and control demands (gradient three). Moreover, the existence of a core DMN network remains controversial since MT and FT subsystems show interdigitated connectivity in these regions (Braga & Buckner, 2017): core DMN may reflect inadequate spatial resolution and/or the integration of informational states across the other subsystems.

This study delineates the role of DMN subsystems in semantic cognition, examining effects of perceptual coupling versus decoupling (in a comparison of reading and autobiographical memory retrieval), modality (words/pictures), abstractness, feature type (valance/spatial) and control demands. We characterise local activation and deactivation, since both are relevant to cognition: connectivity with control regions can increase in demanding semantic decisions even as DMN deactivates (Krieger-Redwood et al., 2016) and multivariate analysis reveals semantic goals are represented in “task-negative” regions (Wang et al., 2021). Deactivation may benefit cognition by suppressing inputs that are irrelevant to the task or context (Amedi et al., 2005; Azulay et al., 2009). Therefore, we examine activating and deactivating voxels within each subsystem for each participant.

## Materials and Methods

The present study investigated DMN activity in 5 independent published fMRI datasets. The key materials and methods are described below but additional details about each dataset are available in previous publications (Study 1: Zhang et al., 2022; Study 2: Krieger-Redwood et al., 2015; Study 3: Hoffman, Binney, & Lambon Ralph, 2015; Study 4: Lanzoni et al., 2020; Study 5: Shao et al., 2022).

### Participants

The samples included: 29 participants (Study 1: mean age ± SD = 20.14 ± 1.26 years, 6 males), 22 participants (Study 2: 23 participants recruited, one removed due to low accuracy, mean age = 23.2 ± 2.9 years, 16 males), 19 participants (Study 3: 20 participants recruited, one removed due to image artefacts, mean age = 25 years, 11 males), 26 participants (Study 4: 27 participants recruited, one excluded due to no behavioural responses being recorded, mean age = 21.5 ± 2.9 years, 9 males) and 176 participants (Study 5: 207 participants recruited, 31 excluded: 25 with missing behavioural data, two with missing or incorrectly recorded imaging data, and four during pre-processing because they exceeded .3mm motion, 20% invalid scans and/or z>2 mean global signal change, mean age = 20.57, 62 males). All participants were right-handed native English speakers, and had normal or corrected-to-normal vision. None had any history of neurological impairment, diagnosis of learning difficulty or psychiatric illness. All participants provided written informed consent prior to taking part. All studies were approved by the local ethics committee.

### Task Procedure

#### Study 1: Reading/autobiographical memory task

The first study compared DMN recruitment during tasks that involved conceptual access driven by visual inputs (in reading comprehension) and internally directed memory retrieval (in autobiographical memory recall). Testing occurred across two consecutive days (See Figure 1A for task design). On Day 1, participants generated their own personal memories from cue words (i.e., Party) outside the scanner. They were asked to identify specific events that they were personally involved in and to provide as much detail about these events as they could, including when and where the event took place, who was involved, what happened, and the duration. They typed these details into a spreadsheet to ensure comparable information was recorded for the different cue words.

**Figure 1.**
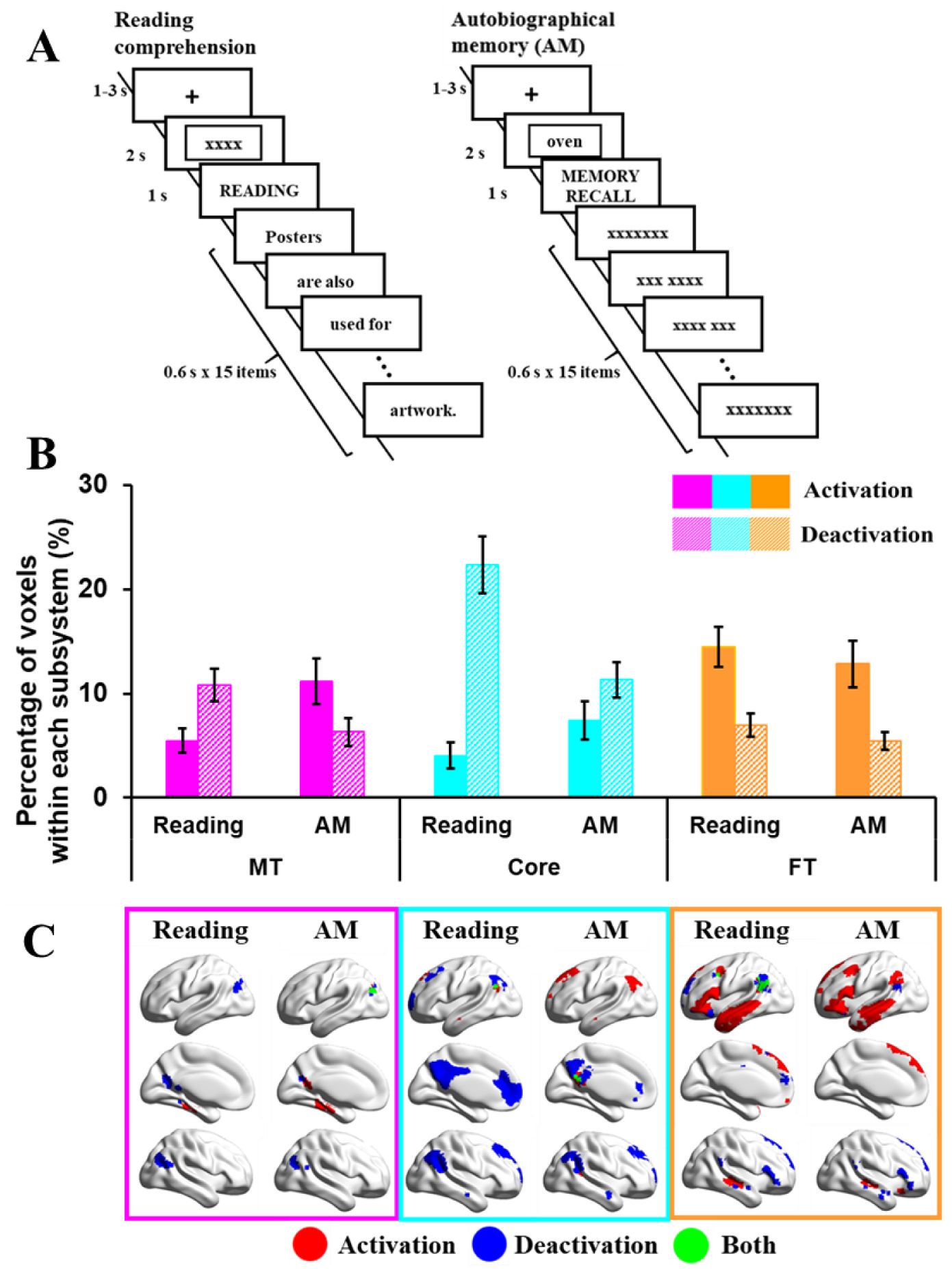
**Panel A** shows the example of task procedures of reading or autobiographical memory (recall) tasks (Study 1). **Panel B** shows the percentage of activating or deactivating voxels in reading or autobiographical memory (recall) tasks. The three colours represent the percentage of voxels extracted from three subsystems (pink = medial temporal regions, cyan = core regions, orange = fronto-temporal regions). Solid bars represent the percentages of activation voxels, and grid bars represent the percentage of deactivation voxels. Error bars represent one standard error. **Panel C** shows the regions of activating voxels (in red colour), deactivating voxels (in blue colour), and the overlapping regions of both activating and deactivating voxels (i.e., regions showed activation in several participants, and deactivation in another group of participants, in green colour).

On the following day, participants were asked to read sentences for comprehension, or to recall their generated personal memories inside the scanner. In reading trials, sentences were presented word by word, after either (1) an autobiographical memory cue word (e.g., Party), creating conflict between reading and personal memory retrieval, or (2) a letter string (e.g., XXX) allowing reading to take place in the absence of conflict from autobiographical memory. We controlled the duration of the sentences by presenting the words on 15 successive slides, combining short words or articles and conjunctions together with nouns on a single slide. In memory recall trials, participants were asked to recall autobiographical memories during the presentation of either (1) meaningful yet unrelated sentences, creating conflict from task-irrelevant patterns of semantic retrieval or (2) letter strings (XXX) allowing autobiographical memory to take place without distracting semantic input. A baseline condition involved presenting meaningless letter strings (i.e., xxxxx) in the absence of a task.

As shown in Figure 1A, each trial started with a fixation cross (1-3s) in the centre of the screen. Then either an autobiographical memory cue word or a letter string appeared for 2s. During the presentation of the cue word, participants were asked to recall their personal memories related to this item. Next, the task instruction (i.e., READING or MEMORY RECALL) was presented for 1s. Following that, words from sentences or letter strings were presented, with each one lasting 600ms. On memory recall trials, participants were asked to keep thinking about their autobiographical memory, in as much detail as possible, until the end of the trial.

#### Study 2: Word/picture semantic judgement task

The second study compared DMN recruitment across semantic tasks involving words and pictures and manipulated the difficulty of these decisions by contrasting strong and weak associations. A three alternative forced choice (3AFC) format was used for all conditions (see Figure 2A for example stimuli and task design). The verbal task involved auditory presentation of a probe word, and response options presented as written words. The picture task used photographs of the probes, targets and distracters.

**Figure 2.**
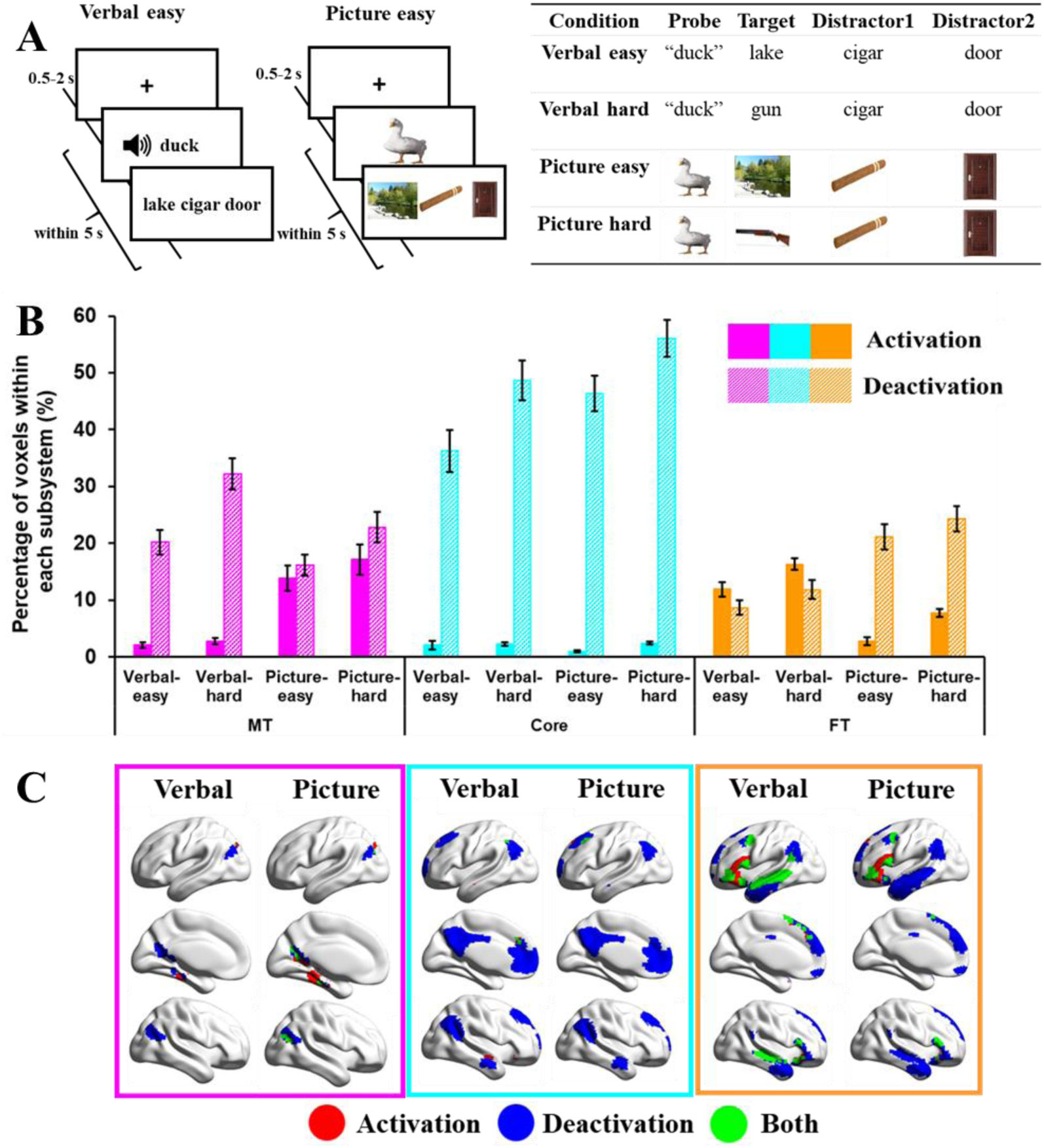
**Panel A** shows the example of task procedures of word or picture matching tasks (Study 2). **Panel B** shows the percentage of activating or deactivating voxels in verbal or picture matching tasks. The three colours represent the percentage of voxels extracted from three subsystems (pink = medial temporal regions, cyan = core regions, orange = fronto-temporal regions). Solid bars represent the percentages of activation voxels, and grid bars represent the percentage of deactivation voxels. Error bars represent one standard error. **Panel C** shows the regions of activating voxels (in red colour), deactivating voxels (in blue colour), and the overlapping regions of both activating and deactivating voxels (i.e., regions showed activation in several participants, and deactivation in another group of participants, in green colour).

Participants made easy and hard associative judgements: they were presented with a spoken word or picture probe, together with three word/picture response options on the screen and instructed to select the item most strongly related to the probe. The probes and targets either shared a strong association (for easy trials), or a weak association (for more difficult trials). For example, an easy association might involve the probe “duck”, and three answer choices such as lake–cigar–door. A harder trial would require participants to link “duck” with gun – an association that is less frequently encountered. Strong associations are thought to be retrieved relatively automatically, since the probe establishes a context that strongly anticipates the target; in contrast, for weak associations, control processes are needed to focus retrieval on non-dominant semantic features that are relevant to the linking context (e.g., a duck can be hunted; a gun is used for hunting; Davey et al., 2016; Jefferies, 2013; Lambon Ralph et al., 2017).

As shown in Figure 2A, each trial started with a fixation screen for a jittered interval (500–2000 ms) followed by the trial (probe and options). Participants were required to make a response, which triggered the next trial; if no response was given after 5 s, the experiment moved onto the next trial.

#### Study 3: Abstract/concrete word synonym judgement task

The third study examined DMN responses to abstract and concrete concepts in a synonym judgement task (see Figure 3A for task design, and Supplementary Table S5 for example stimuli). This study consisted of 200 trials (100 concrete and 100 abstract words); imageability was significantly higher for concrete than abstract words (t = 82, p < .001). On each trial, participants were presented with a written probe word with three choices below it (a semantically related target and two unrelated foils), and the probe word, target word and two distractors had similar imageability ratings (i.e., abstractness) within each trial. They were asked to select the word that was most similar in meaning to the probe. Prior to each decision, participants were presented with a written cue consisting of two short sentences. On half of the trials, the cue ended with the probe word and placed it in a meaningful context (contextual cue condition). On the remaining trials, the cue did not contain the probe and was not related in meaning to the subsequent judgement (irrelevant cue condition). Participants were unaware when reading the cue whether it would be helpful for their next decision.

**Figure 3.**
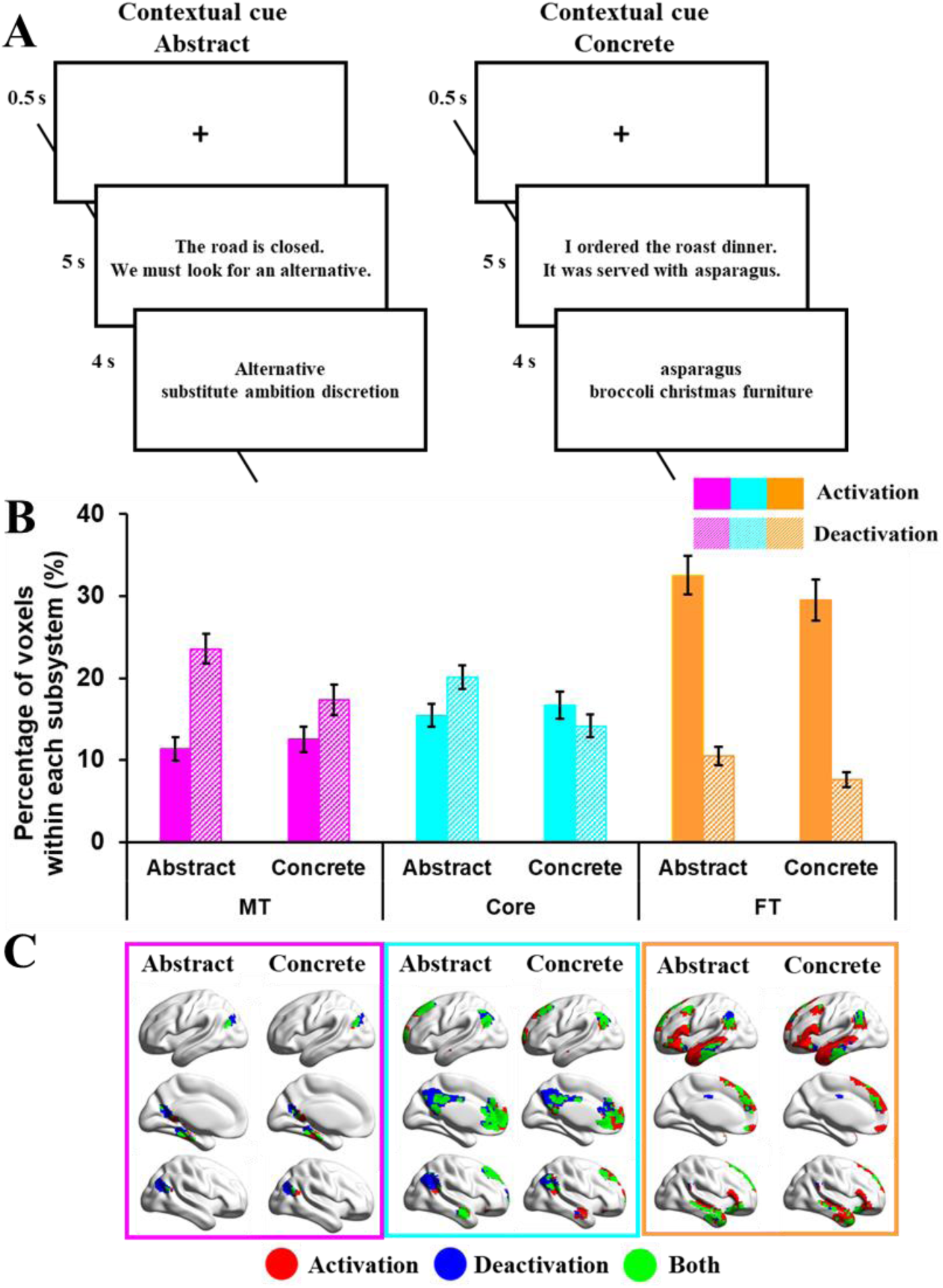
**Panel A** shows the example of task procedures of abstract or concrete tasks (Study 3). **Panel B** shows the percentage of activating or deactivating voxels in abstract or concrete tasks. The three colours represent the percentage of voxels extracted from three subsystems (pink = medial temporal regions, cyan = core regions, orange = fronto-temporal regions). Solid bars represent the percentages of activation voxels, and grid bars represent the percentage of deactivation voxels. Error bars represent one standard error. **Panel C** shows the regions of activating voxels (in red colour), deactivating voxels (in blue colour), and the overlapping regions of both activating and deactivating voxels (i.e., regions showed activation in several participants, and deactivation in another group of participants, in green colour).

As shown in Figure 3A, each trial began with a fixation cross presented in the centre of the screen for 500 ms, followed by the cue. Participants were instructed to read the cue carefully and to press a button on the response box when they had finished reading. The cue remained on screen for 5 s. The judgement probe and three choices were then presented and participants responded by pressing one of three buttons on a response box held in their right hand. The stimuli remained on screen for 4 s, at which point the next trial began. Stimuli were presented in blocks of two trials (total duration = 19 s) with the two trials in each block being taken from the same experimental condition. There were 150 blocks in total and blocks from different conditions were presented in a pseudo-random order. A fixation block of 19 s, in which no stimuli were presented, occurred after every five blocks of task.

#### Study 4: Emotional/spatial cues task

The fourth study examined semantic judgements about ambiguous words preceded by portrayals of facial emotions and spatial locations (see Figure 4A for task design and example stimuli). English homonyms with more than one meaning were selected as stimuli; they had different interpretations associated with different facial expressions (e.g. JAM with TRAFFIC is associated with frustration while JAM with STRAWBERRY is associated with pleasure) but also different locations (e.g. MOTORWAY for TRAFFIC JAM and SUPERMARKET for STRAWBERRY JAM). Four target words were generated for each probe, two for each interpretation. For instance, the probe jam appeared in four trials, twice paired with a target referring to traffic (jam-horn or jam-delay) and twice paired with a target referring to the alternative interpretation (jam-spoon or jam-bread).

**Figure 4.**
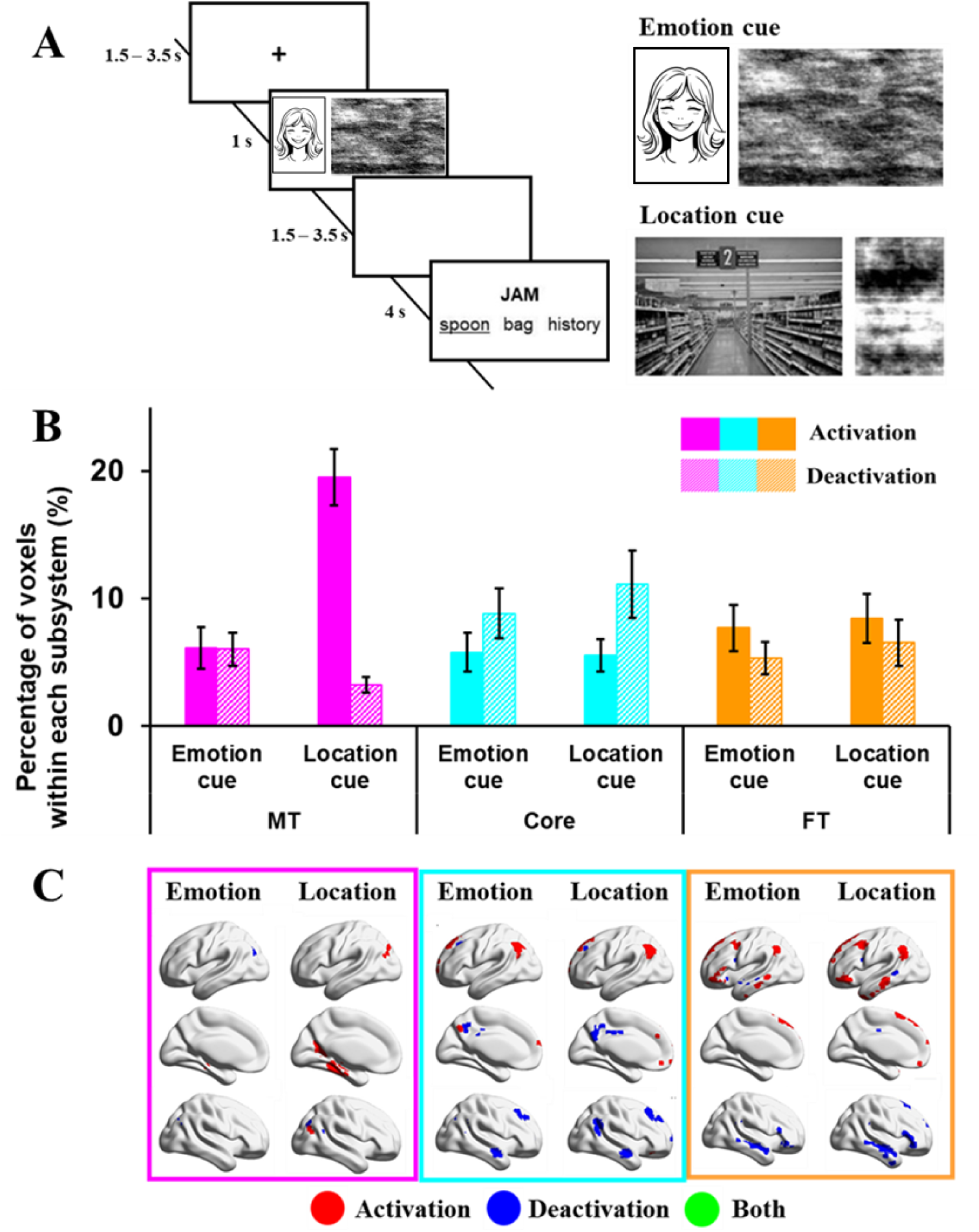
**Panel A** shows the example of task procedures of emotional or spatial cueing tasks (Study 4). **Panel B** shows the percentage of activating or deactivating voxels in emotional or spatial cueing tasks. The three colours represent the percentage of voxels extracted from three subsystems (pink = medial temporal regions, cyan = core regions, orange = fronto-temporal regions). Solid bars represent the percentages of activation voxels, and grid bars represent the percentage of deactivation voxels. Error bars represent one standard error. **Panel C** shows the regions of activating voxels (in red colour), deactivating voxels (in blue colour), and the overlapping regions of both activating and deactivating voxels (i.e., regions showed activation in several participants, and deactivation in another group of participants, in green colour).

Pictures of facial emotional expressions and spatial locations were used to prime the relevant meaning of the homonym. Each picture was used only once across the experiment, so that participants could not predict the subsequent probe word from the cue. Images of facial expressions were chosen from the Radboud Faces Database (Langner et al., 2010) and included eight basic emotions: happy, angry, sad, disgusted, contemptuous, surprised, neutral, fearful. Pictures of spatial contexts were downloaded from Google images.

The experiment included 2-cue (emotion *and* location), 1-cue (either emotion *or* location) and no-cue (scrambled images) trials. The emotion and location cues were presented simultaneously in the 2-cue condition, while for the 1-cue conditions, images were paired with a meaningless scrambled image. The position of the emotion and location cues on the screen (on the left or right-hand side) was counterbalanced within each run. In the no-cue condition, two scrambled images were presented. We used this study to clarify the effect of emotion in driving DMN function; as such, only the 1-cue condition was relevant because each trial contained either an emotion *or* location cue, whereas the 2-cue condition contained both an emotion *and* a location cue for each trial (therefore, the 2-cue condition does not separate the two dimensions).

As shown in Figure 4A, each trial began with a fixation cross (1500–3000 ms) followed by cue images for 1000 ms, and then a blank screen (1500–3000 ms). Following this, a probe word was presented above a target word and two unrelated distracters, triggering the onset of the decision-making period. The probe and choices remained visible for a fixed interval of 4000 ms. The assignment of the emotion-related and location-related distractors to the different conditions was counterbalanced within participants, such that each probe appeared twice with an emotion-related distractor and twice with a location-related distractor.

#### Study 5: Resting-state scan

Participants took part in a 9-minute resting-state fMRI scan. They were instructed to focus on a fixation cross with their eyes open, and not to think about anything in particular. A structural scan was also obtained in the same session.

### Analysis of the percentage of activated/deactivated voxels

To understand the role of subdivisions of DMN in semantic tasks, we extracted numbers of activating and deactivating voxels, relative to the implicit baseline, for each participant within three DMN subsystems – medial temporal (MT), core (Core) and fronto-temporal (FT) – across the four fMRI datasets described above, which contrasted internally-oriented (recall) and externally-oriented (reading) states (Study 1), input modalities (picture/verbal; Study 2), the concreteness of words (abstract/concrete; Study 3) and the nature of cues in semantic decisions (emotion/location; Study 4). The implicit baseline consisted of periods of unmodelled time, for example, between blocks of trials, in which participants were not instructed to do anything. Although “resting periods” are not an ideal baseline for semantic activation (and deactivation), given that semantic cognition will continue (Binder et al., 1999), here this approach provided an opportunity to compare responses across studies to relatively common baseline. The networks of interest were defined using the Yeo et al. 17-network parcellation of 1000 resting-state fMRI datasets projected into individual space (Yeo et al., 2011).

In the analysis presented in the main body, individual activation maps were thresholded at z=1.96 and activating/deactivating voxels were represented as a percentage of voxels in each subsystem. Confirmatory analyses using different thresholds (z=2.3; z=2.6) are presented in Supplementary Materials and showed similar effects. The advantage of this method is that it can reveal brain areas in which voxels both activate and deactivate in response to task demands; this pattern might correspond to a more specific response or connection pattern during a task which cannot be identified if activation levels are averaged across all the voxels within a region. The more standard univariate approach, averaging across voxels, is reported in Supplementary Materials for comparison.

Repeated-measures ANOVAs examined interactions of task by network for activated and deactivated voxels separately. These were followed by analyses of each network separately. When necessary, we also report *post-hoc* Bonferroni-corrected t-tests to characterise task interactions within networks. We also identified the typical location of activating and deactivating voxels by adding together the individual-level maps of all conditions across four studies and thresholding these combined maps at 20% percent of all datasets. All brain figures were created using BrainNet Viewer (http://www.nitrc.org/projects/bnv/; Xia et al., 2013).

### Functional connectivity of the commonly activated and deactivated DMN regions

In Study 5, we examined the intrinsic connectivity of the commonly activated and deactivated regions for each of the three subsystems (i.e., six seeds in total) in resting-state fMRI. In a first-level analysis, we computed whole-brain seed-to-voxel correlations for each seed in the same model after the BOLD time series were pre-processed and denoised. For the group-level analysis, we extracted seed-to-voxel functional connectivity at rest for 176 participants and performed contrasts between the functional connectivity maps from activating and deactivating seeds across the different subsystems (i.e., MT versus Core, FT versus Core, MT versus FT). Group-level analyses in CONN used a voxel threshold of p < 0.001 and were cluster-size FWE corrected at p < .05 (two-sided tests, Bonferroni corrected to p < .017 to account for the three contrasts). The functional connectivity maps for each contrast were uploaded to Neurovault (https://neurovault.org/collections/14749/; Gorgolewski et al., 2015) and decoded using Neurosynth (Yarkoni et al., 2011). The top 20 terms relevant to the maps were rendered as word clouds using a word cloud generator (https://www.wordclouds.com/).

### Gradient analysis of DMN connectivity patterns

To locate the intrinsic connectivity of the commonly activated and deactivated regions for each DMN subsystem within the topographical space defined by whole-brain cortical gradients, we computed the spatial correlations between our intrinsic connectivity maps and the top three cortical gradients defined by Margulies et al. (2016). The cortical gradients were extracted using diffusion embedding from a whole-brain connectivity matrix and represented components of spatial variance in intrinsic connectivity across the cortical surface (maps of Gradient 1 to 3 are from Margulies et al., 2016, https://neurovault.org/collections/1598/). Gradient 1 (the principal gradient, explaining the most variance) differentiates the connectivity patterns of heteromodal DMN (with positive values) and sensory-motor regions (with negative values). Gradient 2 captures the separation in connectivity between visual (positive) and auditory-motor (negative) connectivity patterns. Gradient 3 differentiates heteromodal regions that support more controlled cognition (positive) from core DMN (negative). These spatial correlations with each gradient were computed at the individual level, allowing us to perform repeated-measures ANOVAs that examined the location of activating and deactivating voxels for each DMN subsystem in gradient space.

## Results

### Analysis of the percentage of activated/deactivated voxels

#### Study 1: Reading/autobiographical memory task

We extracted the percentage of activating and deactivating voxels during *reading* and autobiographical *recall* within each subsystem (see Figure 1). The conflict and no conflict conditions did not show any differences, so here we report repeated-measures ANOVAs including task (*reading* or *recall*) by DMN subsystem (MT, Core or FT) for activated or deactivated voxels (models that include the conflict manipulation are reported in Supplementary Materials).

##### Activation

Activating voxels showed a main effect of task (*F*(1, 28) = 5.36, *p* = 0.028, *η^2^* = .16), with stronger activation for *recall* than for *reading*, a significant main effect of DMN subsystem (*F*(2, 56) = 22.14, *p* < .001, *η^2^* = .44), and a significant interaction effect between task and DMN subsystem (*F*(2, 56) = 22.18, *p* < .001, *η^2^* = .44). Paired t-tests (corrected for the number of comparisons) compared the percentage of voxels activating across subsystems: the FT subsystem showed more activation than MT (t(28) = 4.22, Bonferroni-corrected *p* < .001), while the core showed less activation than both MT (t(28) = −2.56, Bonferroni-corrected *p* = .048) and FT subsystems (t(28) = −5.88, Bonferroni-corrected *p* < .001). To understand the interaction between subnetwork and task, paired t-tests compared *reading* and *recall* in each subsystem: in the MT and Core, *reading* elicited less activation than *recall* (MT: t(28) = −3.45, Bonferroni-corrected *p* = 0.006; Core: t(28) = −3.20, Bonferroni-corrected *p* = 0.009), while there was no significant difference in the FT subsystem (Bonferroni-corrected *p* = 0.249).

##### Deactivation

Deactivating voxels showed a significant main effect of task (F(1, 28) = 28.13, *p* < .001, *η^2^* = .50), with more deactivation for *reading* than *recall*, a significant main effect of DMN subsystem (F(2, 56) = 22.01, *p* < .001, *η^2^*= .44), and a significant interaction effect between task and DMN subsystem (F(2, 56) = 21.94, *p* < .001, *η^2^* = .44). The Core subsystem deactivated more than the MT and FT subsystems (Bonferroni-corrected *p* < .001), and there was no significant difference in deactivation between the MT and FT subsystems (Bonferroni-corrected *p* = .297). To understand the interaction effect, paired t-tests were conducted between *reading* and *recall* in each subsystem (see Table 2). *Reading* elicited more deactivation than *recall* in the MT and Core subsystems (MT: t(28) = 4.18, Bonferroni-corrected *p* < .001; Core: t(28) = 5.57, Bonferroni-corrected *p* < .001), and there was no significant difference in the FT (FT: t(28) = 2.11, Bonferroni-corrected *p* = .132).

###### Study 1 Summary

The Core subsystem showed substantial task-related deactivation, especially during reading. The MT subsystem activated more and deactivated less during recall than reading. The FT subsystem was activated during both reading and recall and showed no differences between tasks. These results are consistent with the view that DMN subsystems have unique functional responses. Core DMN appears to be most decoupled from visual and attentional states (Zhang et al., 2022), even for tasks that involve semantic cognition and that are relatively naturalistic (i.e., reading as opposed to semantic decisions). The MT subsystem, in contrast, shows stronger activation to memory-based tasks, while the FT subsystem activates to both externally oriented (reading) and internally oriented (recall) tasks, consistent with the view that this DMN subsystem supports semantic cognition (Andrews-Hanna and Grilli, 2021). However, reading is a highly verbal task, while autobiographical memory is likely to involve more visuo-spatial processes that support internal scene construction (Andrews-Hanna et al., 2014; Andrews-Hanna and Grilli, 2021). To establish whether differences between the FT and MT subsystems reflect perceptually coupled versus decoupled aspects of cognition, or alternatively might reflect the recruitment of more abstract/verbal versus visuospatial codes, we compared the activation and deactivation of these subsystems across externally presented verbal and picture-based semantic decisions in Study 2.

#### Study 2: Word/picture semantic judgement task

This task allowed us to compare activation and deactivation for semantic decisions involving words and pictures, which also varied in difficulty. The probe-target pairs were either strongly associated (easy trials) or weakly associated (hard trials). We extracted the percentage of activating and deactivating voxels in these four conditions: word-easy, word-hard, picture-easy, picture-hard, within each subsystem, and performed a 2 (modality: word or picture) by 2 (difficulty: easy or hard) by 3 (DMN subsystem: MT, Core or FT) repeated-measures ANOVA for activating and deactivating voxels, respectively.

##### Activation

For activating voxels, there was no main effect of modality (*F*(1, 21) = 2.33, *p* = 0.14, *η^2^* = .10), a significant main effect of difficulty (*F*(1, 21) = 15.50, *p* < .001, *η^2^*= .43), with hard conditions activating more voxels than easy conditions, and a significant main effect of DMN subsystem (*F*(1, 21) = 33.31, *p* < .001, *η^2^* = .61), with the Core subsystem showing fewer activating voxels relative to MT (t(21) = −5.66, Bonferroni-corrected *p* < .001) or FT (t(21) = −14.61, Bonferroni-corrected *p* < .001), and no significant difference between the MT and FT (t(21) = .60, Bonferroni-corrected *p* > 1). There were interaction effects between task modality and DMN subsystem (*F*(1, 21) = 73.32, *p* < .001, *η^2^* = .78), and between difficulty and DMN subsystem (*F*(1, 21) = 10.63, *p* < .001, *η^2^* = .34), but no other significant interactions.

To understand the significant two-way interactions for activation, we conducted 2 (modality: word or picture) by 2 (difficulty: easy or hard) repeated-measures ANOVAs for each subsystem. In MT, activating voxels showed a significant main effect of task modality (*F*(1, 21) = 38.33, *p* < .001, *η^2^* = .65), with picture conditions eliciting more activating voxels than word conditions; there was no main effect of difficulty (*F*(1, 21) = 3.33, *p* =.08, *η^2^* = .14) and no interaction effect between modality and difficulty (*F*(1, 21) = 2.26, *p* = .15, *η^2^* = .10). In the core subsystem, activating voxels showed no main effect of modality (*F*(1, 21) = 1.09, *p* = .31, *η^2^* = .05), a near significant main effect of difficulty (*F*(1, 21) = 3.91, *p* =.061, *η^2^* = .16) with hard conditions activating marginally more voxels than easy conditions, and no significant interaction effect (*F*(1, 21) = 3.06, *p* = .095, *η^2^* = .13). In the FT, activating voxels showed a main effect of modality (*F*(1, 21) = 119.29, *p* < .001, *η^2^* = .85), with verbal conditions eliciting more activation than pictures, a significant main effect of difficulty (*F*(1, 21) = 35.69, *p* < .001, *η^2^* = .63), with hard conditions activating more voxels than easy conditions, and no significant interaction effect (*F*(1, 21) = .10, *p* = 0.76, *η^2^* = .005).

##### Deactivation

Deactivating voxels showed a main effect of modality (*F*(1, 21) = 4.70, *p* = .042, *η^2^* = .18), with picture conditions eliciting more deactivation than word conditions, and a significant main effect of difficulty (*F*(1, 21) = 25.73, *p* < .001, *η^2^* = .55), with hard conditions eliciting more deactivation than easy conditions. There was also a main effect of DMN subsystem (*F*(1, 21) = 122.75, *p* < .001, *η^2^* = .85), with the Core showing more deactivation relative to MT (t(21) = 11.05, Bonferroni-corrected *p* < .001) or FT (t(21) = 14.19, Bonferroni-corrected *p* < .001), and MT showing more deactivation relative to FT (t(21) = 3.54, Bonferroni-corrected *p* = . 006). There were interactions between modality and subsystem (*F*(1, 21) = 33.14, *p* < .001, *η^2^* = .61) and difficulty and subsystem (*F*(1, 21) = 9.18, *p* < .001, *η^2^* = .30), but no other significant interactions.

To understand the significant two-way interactions for deactivation, we conducted 2 (modality: word or picture) by 2 (difficulty: easy or hard) repeated-measures ANOVAs for each subsystem. MT showed a main effect of modality (*F*(1, 21) = 6.43, *p* = .019, *η^2^* = .23), with words eliciting more deactivation than pictures, a main effect of difficulty (*F*(1, 21) = 28.49, *p* < .001, *η^2^* = .58), with hard conditions eliciting more deactivation than easy conditions, and no significant interaction (*F*(1, 21) = 1.90, *p* = 0.18, *η^2^* = .08). Core DMN showed a main effect of task modality (*F*(1, 21) = 8.26, *p* = .009, *η^2^*= .28), with pictures eliciting more deactivation than words, a main effect of difficulty (*F*(1, 21) = 20.30, *p* < .001, *η^2^* = .49), with more deactivation for hard than easy conditions, and no significant interaction (*F*(1, 21) = .37, *p* = 0.55, *η^2^* = .02). FT showed the opposite effect of modality (*F*(1, 21) = 31.33, *p* < .001, *η^2^* = .60), with more deactivation for pictures than for words, a main effect of difficulty (*F*(1, 21) = 5.37, *p* = .031, *η^2^* = .20), with more deactivation for hard than easy conditions, and no interaction (*F*(1, 21) = .00, *p* = .99, *η^2^* = .00).

###### Study 2 Summary

The MT subsystem showed more activation for pictures and more deactivation for words, while FT showed the opposite pattern. Core DMN showed little activation for either words or pictures, yet more deactivation for pictures, broadly consistent with the proposal that this network shows perceptual decoupling (Andrews-Hanna, Smallwood, and Spreng, 2014). The DMN subsystems also showed different responses to the difficulty manipulation: while all three networks showed more deactivation in response to harder judgements, in line with the ‘task-negative’ expectation for DMN regions (Greicius et al., 2003; Raichle et al., 2001), the FT subsystem also showed significantly more activation for difficult semantic trials. FT shows *both* activation and deactivation in response to harder semantic decisions, suggesting it shows more specific patterns of response under demanding circumstances, a pattern that we refer to as “tuning”.

These results help to constrain interpretations of functional distinctions between DMN subsystems. We found that MT is not only implicated in autobiographical memory but also in external tasks involving pictures as opposed to words, suggesting it supports visuo-spatial representations across both external and internal modes of cognition. In contrast, the FT subsystem is more strongly implicated in verbal semantic tasks, suggesting that this network might support abstract aspects of semantic processing. The next study tests this proposal directly by comparing responses to semantic decisions involving concrete and abstract words.

#### Study 3: Abstract/concrete word synonym judgement task

We extracted the percentage of activating and deactivating voxels during semantic decisions involving abstract and concrete words within each subsystem and performed a 2 (task: abstract or concrete) by 3 (DMN subsystem: MT, Core or FT) repeated-measures ANOVA examining activation and deactivation voxels, respectively. There were no significant effects of cueing, therefore this experimental factor is omitted below (see Supplementary Materials for full analysis).

##### Activation

For activating voxels, there was no main effect of concreteness (*F*(1, 18) = .10, *p* = .75, *η^2^* = .006), a significant main effect of DMN (*F*(2, 36) = 30.02, *p* < .001, *η^2^* = .63), and a significant interaction effect between concreteness and DMN subsystem (*F*(2, 36) = 13.45, *p* < .001, *η^2^* = .43). FT activated more than MT (t(18) = 5.66, Bonferroni-corrected *p* < .001) and Core (t(18) = 6.80, Bonferroni-corrected *p* < .001), while the difference in activation between MT and Core was not significant (t(18) = 2.08, Bonferroni-corrected *p* = .159). To understand the interaction effect, paired t-tests were conducted between abstract and concrete words in each subsystem: there was no effect of this manipulation in either the MT or Core subsystems (MT: t(18) = 1.78, Bonferroni-corrected *p* = .276; Core: t(18) = 1.22, Bonferroni-corrected *p* = .711), while abstract words elicited more activation than concrete words in the FT subsystem (t(18) = 3.80, Bonferroni-corrected *p* = .003).

##### Deactivation

For deactivating voxels, there were main effects of concreteness (*F*(1, 18) = 59.94, *p* < .001, *η^2^* = .77), and DMN subsystem (*F*(2, 36) = 20.73, *p* < .001, *η^2^* = .54), plus an interaction between them (*F*(2, 36) = 5.86, *p* = .018, *η^2^* = .25. FT deactivated less than MT (t(18) = 5.40, Bonferroni-corrected *p* < .001) and Core subsystems (t(18) = 6.79, Bonferroni-corrected *p* < .001), while the MT and the Core subsystems did not differ (t(18) = 1.63, Bonferroni-corrected *p* = .36). Paired t-tests to examine the interaction showed that abstract words elicited more deactivation than concrete words in all three subsystems (MT: t(18) = 4.68, Bonferroni-corrected *p* < 0.001; Core: t(18) = 9.17, Bonferroni-corrected *p* < 0.001; FT: t(18) = 5.54, Bonferroni-corrected *p* < 0.001), but this effect was largest in the Core.

###### Study 3 Summary

The FT subsystem showed stronger activation to *abstract* than *concrete* concepts, consistent with the interpretation that it supports abstract aspects of meaning. FT DMN regions, for example, in lateral anterior temporal lobes, are thought to lie at the end of a processing stream that supports the abstraction of meaning from diverse sensory-motor features. Yet the FT subsystem also showed greater deactivation for abstract concepts (i.e., a tuning effect, like the effect of difficulty found for FT in Study 2). Abstract concepts along with more difficult semantic decisions might require a more selective pattern of DMN activation and connectivity. The MT subsystem was equally activated by *abstract* and *concrete* conditions, and both MT and Core DMN deactivated more to *abstract* words. Core DMN also appeared to show more activating voxels in this task, in which verbal semantic decisions were made following a meaningful sentence cue, as opposed to Study 2, in which semantic decisions occurred in the absence of a context.

The evidence presented so far demonstrates greater activation for verbal semantic tasks and abstract concepts in FT DMN, and for picture semantic tasks and autobiographical memory retrieval in MT DMN, although we also found that FT DMN responds in a similar way to internally-oriented and externally-oriented tasks that involve meaning (Study 1). Abstract words also entail affective processing to a greater extent than concrete concepts (Kousta et al., 2011; Vigliocco et al., 2014). To clarify the effect of emotion in driving differences in DMN function, the next study compared activation for facial portrayals of emotion and pictures of spatial locations in the three subsystems.

#### Study 4: Emotional/spatial cues task

We extracted the percentage of activating and deactivating voxels within each subsystem during semantic decisions that followed emotion cues (facial portrayals of emotion) and visuo-spatial cues (photographs of scenes) and performed a 2 (cue type: emotion or location) by 3 (DMN subsystem: MT, Core or FT) repeated-measures ANOVA for the percentage of activated and deactivated voxels, respectively. An additional analysis examined the effect of multiple cues on DMN recruitment (the 2-cue condition that leveraged both location and emotion cues within a trial; since this analysis does not reveal effects of emotion versus scenes, these results are reported in Supplementary Materials).

##### Activation

Activation showed a main effect of cue type (*F*(1, 25) = 7.64, *p* = .011, *η^2^* = .23) with spatial cues activating more voxels than emotional cues, a main effect of DMN (*F*(2, 50) = 13.24, *p* < .001, *η^2^* = .35), and a significant interaction effect between cue type and DMN subsystem (*F*(2, 50) = 26.76, *p* < .001, *η^2^* = .52). The MT and FT activated more than the Core (t(25) = 4.39, Bonferroni-corrected *p* < .001 for the MT, t(25) = 3.35, Bonferroni-corrected *p* = .009 for the FT), while the MT and FT subsystems did not differ (t(25) = 1.96, Bonferroni-corrected *p* = .183). To understand the interaction effect, paired t-tests compared the response to emotion and spatial cues in each subsystem: emotional cues elicited less activation than spatial cues in MT (t(25) = 5.07, Bonferroni-corrected *p* < 0.001), while there was no significant difference either in the Core or FT subsystems (Bonferroni-corrected *p* > 1).

##### Deactivation

Deactivating voxels showed no main effect of cue type (*F*(1, 25) = .025, *p* = .88, *η^2^* = .001), a main effect of DMN subsystem (*F*(2, 50) = 9.11, *p* < .001, *η^2^* = .27), and a significant interaction (*F*(2, 50) = 3.90, *p* = .027, *η^2^* = .14). MT and FT deactivated less than Core DMN (t(25) = 4.84, Bonferroni-corrected *p* < .001 for MT, t(25) = 3.46, Bonferroni-corrected *p* = .006 for FT), while the MT and FT subsystems did not differ (t(25) = 1.15, Bonferroni-corrected *p* = .786). Paired t-tests compared deactivation in response to emotion and spatial cues for each subsystem but no significant differences were found (t(25) < 2.07, Bonferroni-corrected *p* > .147).

###### Study 4 summary

The MT subsystem activated significantly more to *spatial* than *emotional* cue conditions. The Core subsystem deactivated to both conditions without a significant difference between them. The FT subsystem activated to both conditions equally, confirming that different kinds of meaningful features can activate this network. These results suggest that even though valence is thought to be a crucial component of the meaning of abstract words (Kousta et al., 2011; Vigliocco et al., 2014), a simple response to valence is unlikely to account for the role of the FT subsystem in abstract semantics. The results also provide a further demonstration that activation in the MT subsystem relates to visual-spatial coding.

### Regions activated and deactivated in DMN subsystems

The subsystems of DMN showed different functional responses across a set of semantic tasks. The FT subsystem showed an increase in the number of activated voxels (relative to the implicit baseline) for both reading and autobiographical memory and showed a stronger response to words and abstract concepts; this suggests it supports semantic tasks that involve abstract concepts across both perceptually coupled and decoupled tasks. The MT subsystem showed an increase in activated voxels (relative to the implicit baseline) for autobiographical memory and pictorial semantic judgements and an increase in deactivated voxels for abstract concepts, suggesting that a visuospatial code is core to its behaviour, although scenes do not need to be generated internally. The core subsystem showed an increased in deactivated voxels (relative to the implicit baseline) in response to most of the externally-oriented tasks but voxels were both activated and deactivated during autobiographical memory, in line with the view that this subsystem supports perceptually decoupled cognition.

We next considered where these regions of activating and deactivating voxels were located within each subsystem across participants and tasks. This is shown in the bottom panels of Figures 1 to 4. The MT subnetwork showed activating voxels, relative to baseline, in medial temporal regions (particularly for autobiographical memory retrieval in Study 1, and across verbal and picture associations in Study 2). The angular gyrus (AG) subcomponent of MT showed more deactivating voxels in Studies 1 and 2, when people accessed non-spatial meanings from inputs, especially in the right hemisphere (in Studies 1 and 3). However, both medial temporal and inferior parietal components of this network contained activating voxels when people made semantic decisions that were cued by images of scenes. Across studies, the MT subsystem reliably showed activating voxels in bilateral medial temporal regions and commonly deactivating voxels in AG and medial occipitoparietal cortex (see Figure 5A).

**Figure 5.**
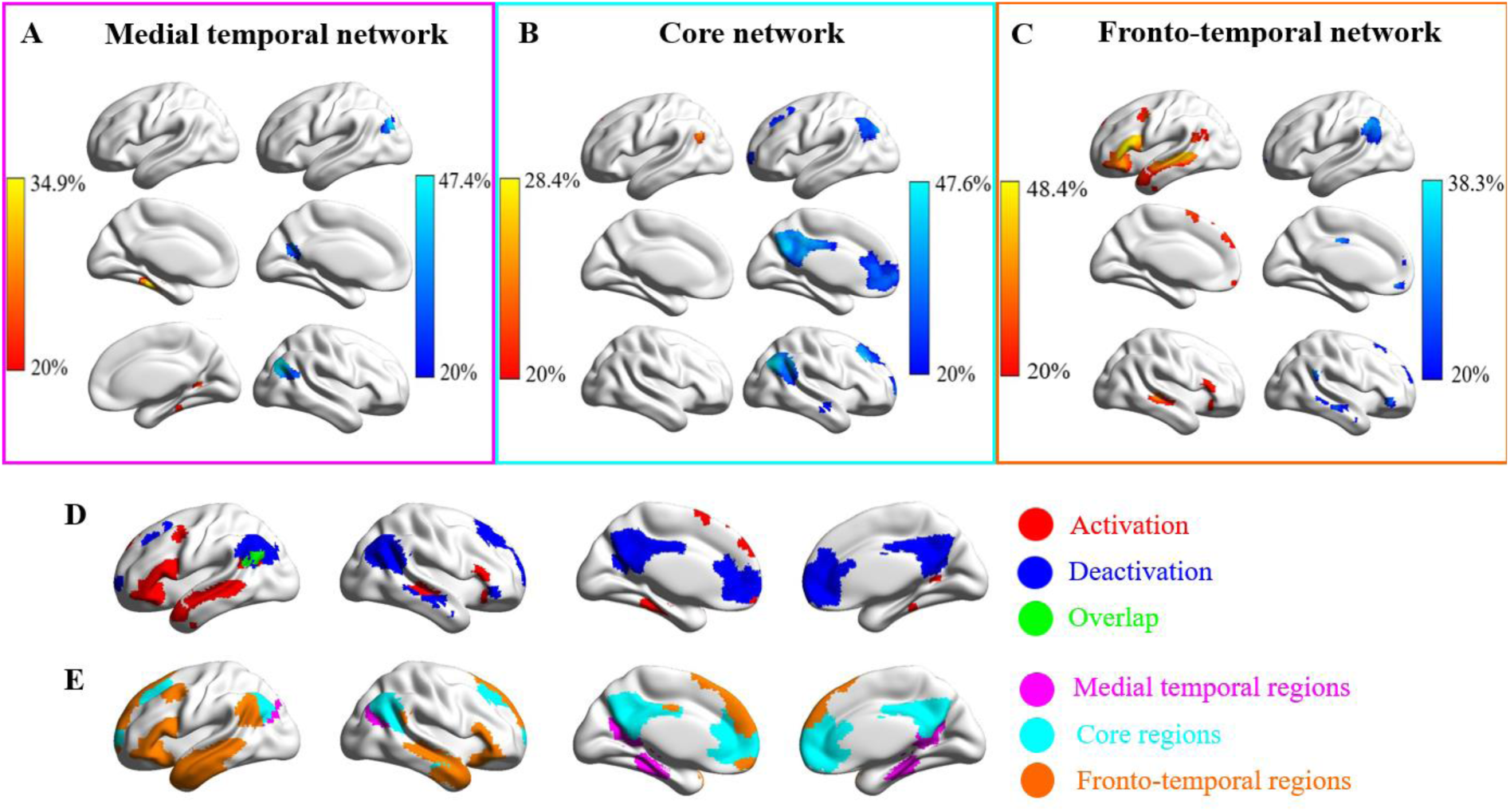
**Panels A-C** show the regions that commonly activated or deactivated in the three subsystems across the four task-fMRI datasets. **Panels A, B, and C** show the regions commonly activated or deactivated in MT, core, and FT subsystems, respectively. The values in the activation or deactivation maps represent the percentage of participants that activated or deactivated in the three subsystems (red = activation, blue = deactivation, green = overlap regions **Panel D** shows the activated and deactivated regions within the whole DMN (binarized from the maps from Panel A-C). **Panel E** shows the maps of three subsystems (pink = medial temporal regions, cyan = core regions, orange = fronto-temporal regions).

**Figure 6.**
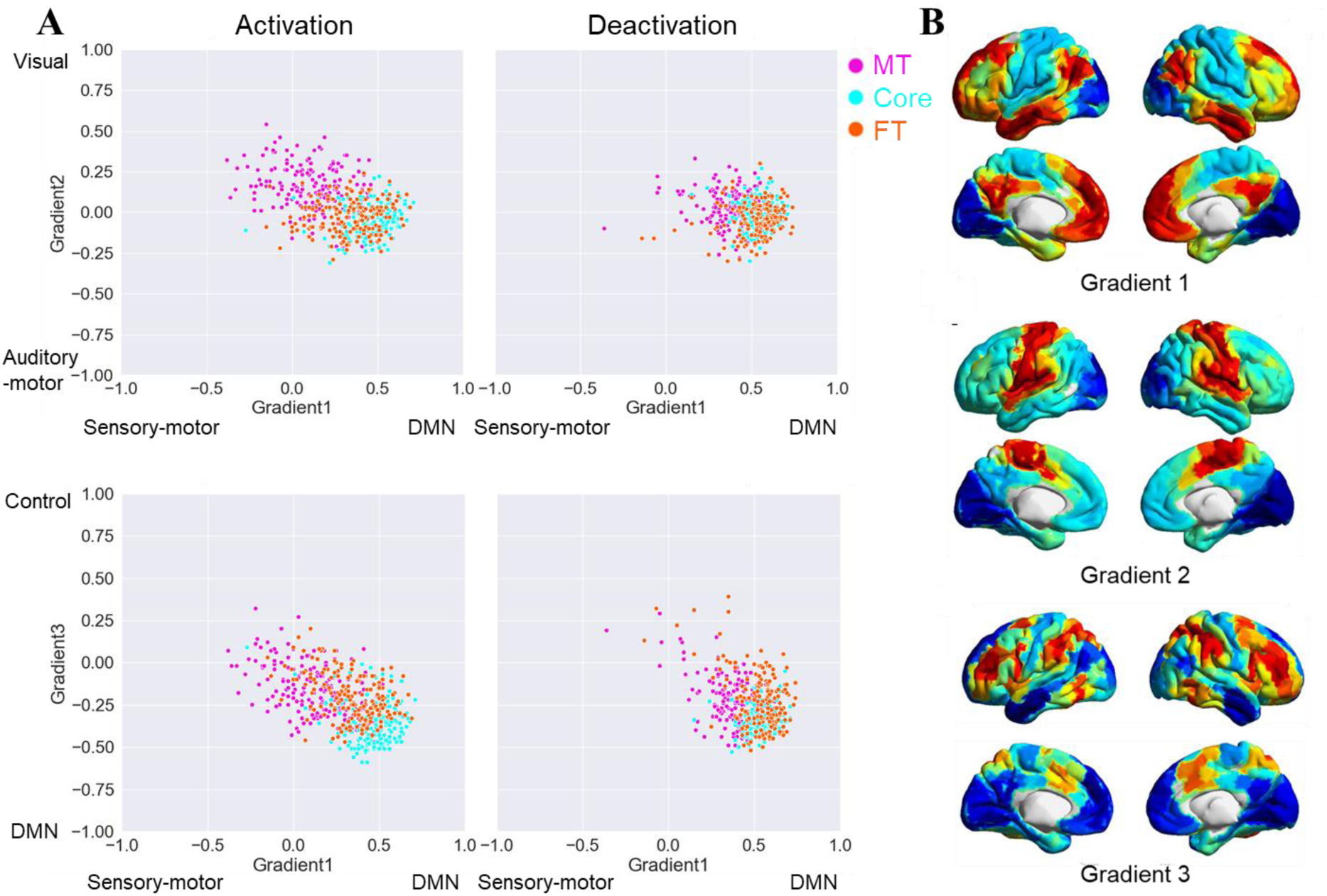
Scatter plots of the individual-level correlations between the functional connectivity maps seeding from commonly activated and deactivated regions in the three subsystems and the three cortical gradients (Figure 6A, Margulies et al., 2016) and the maps of gradient one to three (Figure 6B, taken from Shao et al., 2022).

Core DMN frequently showed deactivation. All nodes of this network showed deactivating voxels during demanding semantic decisions to words and pictures (Study 2), and most showed overlapping areas of activating and deactivating voxels during semantic decisions that followed sentences which were sometimes related in meaning (Study 3). However, activating voxels were seen in left AG when people retrieved autobiographical memories (Study 1) and when they made semantic decisions following previously presented faces and scenes that disambiguated the meaning of words (Study 4). Across studies, the core subsystem showed activating voxels in common areas of left AG, and deactivating voxels in more dorsal and posterior bilateral AG, posterior cingulate cortex (PCC), superior frontal gyrus (SFG) and frontal pole (see Figure 5B).

The FT subsystem showed activating voxels (relative to the implicit baseline) in left hemisphere regions within the semantic network (left inferior frontal gyrus, anterior temporal cortex and angular gyrus) in Studies 1 and 4 involving reading, autobiographical memory and word judgements following face and location cues. In these studies, there were more deactivating voxels in the right hemisphere. For very demanding semantic decisions in Studies 2 and 3, a somewhat different pattern was seen. There were more activating voxels bilaterally (or overlapping areas of activating and deactivating voxels) in inferior frontal gyrus (IFG), and a similar response in anterior temporal cortex across hemispheres (although more deactivation in Study 2 and more activation in Study 3). Across studies, the FT subsystem showed activating voxels in left inferior frontal gyrus (IFG), left AG and bilateral superior temporal gyrus (STG), and deactivating voxels in other regions of left AG as well as right IFG and STG (see Figure 5C).

When combining data across all subsystems (Figure 5), there were reliable regions showing activating voxels across participants and tasks in regions associated with semantic processing, including left temporal cortex, and left inferior frontal gyrus, along with medial temporal cortex. There were common areas containing deactivating voxels in midline anterior and posterior cingulate cortex, broad swathes of right angular gyrus and specific areas of left angular gyrus. Left angular gyrus was the site that most reliably showed both activating and deactivating voxels, which might relate to response ‘tuning’, as specific task-irrelevant representations or connectivity patterns are suppressed (see Studies 2 and 3). In this way, we replicated and extended previous observations that left anterior temporal cortex shows activation in semantic tasks, while angular gyrus deactivates (Humphreys et al., 2015).

### Functional connectivity of the commonly activated and deactivated DMN regions

To understand how areas of activation and deactivation in each DMN subnetwork are functionally connected with the whole brain, we examined functional connectivity at rest. Basic contrasts between the functional connectivity maps for activating and deactivating regions of each subsystem are shown in Supplementary Materials. There was stronger connectivity to visual and motor regions for MT and FT than for Core, consistent with the proposal that the Core DMN is more perceptually decoupled. In cognitive decoding of these connectivity maps, Core DMN was more associated with terms linked to episodic memory and mentalizing, MT was associated with terms such as ‘navigation’, and FT was associated with language terms.

In a final analysis, we situated the functional connectivity maps of commonly activated and deactivated DMN regions in the topographical space defined by overlapping whole-brain cortical gradients (Margulies et al., 2016). Lower spatial correlations with Gradient 1 suggest functional responses that have greater similarity with unimodal regions, while higher correlations suggest more heteromodal and/or abstract responses. Lower spatial correlations with Gradient 2 indicate a stronger motor response, while higher spatial correlations indicate a stronger visual response. Higher spatial correlations with Gradient 3 suggest a response closer to control regions while lower correlations indicate greater similarity with heteromodal regions not associated with control.

For Gradient 1, we found a significant main effect of voxel activation (*F*(1, 175) = 725.63, *p* < .001, *η^2^* = .81), with higher positive (heteromodal/abstract-anchored) correlations for deactivating than activating voxels. There was a significant main effect of DMN subsystem (*F*(2, 350) = 310.91, *p* < .001, *η^2^* = .64); Core DMN showed a higher spatial correlation with Gradient 1 than both MT (Bonferroni-corrected *p* < .001) and FT (Bonferroni-corrected *p* < .001), while FT showed higher correlation than MT (Bonferroni-corrected *p* < .001). There was also a significant interaction between activation and subsystem (*F*(2, 350) = 52.94, *p* < .001, *η^2^* = .23). Gradient 1 correlations were higher for deactivation than for activation for MT (t(175) = 16.85, Bonferroni-corrected *p* < .001), FT (t(175) = 15.50, Bonferroni-corrected *p* < .001) and Core (t(175) = 8.66, Bonferroni-corrected *p* < .001), but this difference between activation and deactivation was smaller for core DMN.

For Gradient 2, we found a main effect of voxel activation (*F*(1, 175) = 42.87, *p* < .001, *η^2^* = .20) with higher spatial correlations (more visual response) seen for regions of activating than deactivating voxels. There was also a main effect of DMN subsystem (*F*(2, 350) = 205.89, *p* < .001, *η^2^* = .54): MT showed higher spatial correlation with the visual end of Gradient 2 than Core DMN (Bonferroni-corrected *p* < .001) or FT (Bonferroni-corrected *p* < .001), consistent with stronger visual representation within this subsystem. There was no difference in Gradient 2 values between the Core and FT (Bonferroni-corrected *p* = 1). Finally, Gradient 2 showed a significant interaction between activation and subsystem (*F*(2, 350) = 96.66, *p* < .001, *η^2^* = .36). Activation regions showed higher correlations with the visual end of Gradient 2 than deactivation regions for MT (t(175) = 11.33, Bonferroni-corrected *p* < .001), while the Core showed the opposite pattern (t(175) = 4.47, Bonferroni-corrected *p* < .001). There was no difference in Gradient 2 correlations across activation and deactivation for FT (t(175) = 1.98, Bonferroni-corrected *p* = .147).

For Gradient 3, there was a main effect of voxel activation (*F*(1, 175) = 13.28, *p* < .001, *η^2^*= .07) reflecting higher correlations with the executive end of this gradient for activating than deactivating voxel regions. There was also a significant main effect of DMN subsystems (*F*(2, 350) = 147.03, *p* < .001, *η^2^* = .46): Core DMN showed lower correlations (less executive response) than MT (Bonferroni-corrected *p* < .001) and FT (Bonferroni-corrected *p* < .001), with no difference between MT and FT (Bonferroni-corrected *p* = .35). Gradient 3 also showed an interaction between activation and subsystem (*F*(2, 350) = 16.94, *p* < .001, *η^2^* = .09). Activating regions showed higher correlations (higher executive response) than deactivating regions for MT (t(175) = 5.71, Bonferroni-corrected *p* < .001), while the Core showed the opposite pattern (less executive response; t(175) = 3.71, Bonferroni-corrected *p* < .001). There was no difference between activating and deactivating voxels within FT (t(175) = 1.75 Bonferroni-corrected *p* = .246).

These results show that deactivating voxels are closer to the heteromodal end of Gradient 1 than activating voxels, especially for the MT and FT subsystems, in line with the view that task-related activation of these subsystems is often driven by sensory inputs. We also found that activating voxels for the MT subsystem were more visual than deactivating voxels on Gradient 2, while the reverse was true of Core DMN (and no difference for FT). This might reflect the key importance of visual codes in the MT subsystem, and perceptual decoupling in Core DMN. Finally, activating voxels in the MT subsystem were closer to the controlled end of Gradient 3 than deactivating voxels, while the reverse was found for Core DMN (again, no difference for FT).

## Discussion

By applying a novel method that examines both activating and deactivating voxels, our study reveals the contribution of DMN subsystems to semantic cognition, and how this is influenced by perceptual decoupling, input modality, abstractness, and spatial versus emotional features. None of the DMN variants were exclusively task negative. Instead, the recruitment of DMN subsystems varied according to the need to maintain information in memory that differs from inputs in the external world, and the requirement to represent visuospatial and abstract conceptual information. Although DMN can be characterized as a unitary whole (Raichle et al., 2001; Yeo et al., 2011), the three subsystems played distinct and complementary roles in semantic cognition which were related to their different locations on a multidimensional space defined by whole-brain gradients of connectivity. Core DMN showed both activating and deactivating voxels during autobiographical memory and when semantic retrieval followed the presentation of earlier information maintained in working memory. It showed almost exclusive voxel deactivation when new trials were presented in the absence of a meaning-based context; this subsystem was also relatively far from unimodal, visual and control regions in gradient space. MT was recruited in internally-oriented tasks involving visual-spatial imagery (autobiographical memory) and by pictorial semantic tasks; it was closer than core DMN to visual and control regions. FT showed more voxel activation for abstract verbal semantic processing and overlapping regions of task activation and deactivation; both activating and deactivating voxels were more numerous during more demanding semantic tasks, suggesting that semantic responses or patterns of cortical connectivity in this subsystem are “tuned” in the face of higher semantic control demands.

Our results align well with a recent topographic account suggesting that the diverse roles of the DMN relate to its spatial location on the cortical mantle (Margulies et al., 2016; Smallwood et al., 2021). In general, this network is distant from sensory-motor cortex on the principal gradient, and this position is suited to cognition that is perceptually-decoupled from the changing external environment or that builds on previously-presented information (for example, when decisions follow sentence cues): we observed this type of response in core DMN. However, this position on the principal gradient also places the DMN at the end of a processing stream (or streams) that allows the integration of features, giving rise to heteromodal representations that support cognition that is both guided by memory and driven by new sensory inputs: we observed this type of response in FT and MT DMN.

The responses we observed for core DMN are broadly in line with a ‘perceptual decoupling’ view of the DMN and not with a ‘task-negative’ view (Greicius et al., 2003; Raichle et al., 2001). Although core DMN showed more deactivation than the MT and FT subsystems across multiple semantic tasks, demonstrating that this subnetwork decouples from visual and attentional states (even during long-term memory retrieval in some circumstances), the numbers of activating and deactivating voxels were more similar when the task involved sustaining meaningful information over time. The ability of core DMN to support conceptual information that is distinct from the unfolding sensory environment might be supported by its greater distance from unimodal and visual ends of the gradients compared with other subnetworks. The functional importance of this distance from visual cortex is also supported by our finding that activating voxels in core DMN were on average more distant from visual cortex than deactivating voxels. Moreover, the observation that core DMN voxels can activate over baseline during the maintenance of contextual information as well as during autobiographical memory retrieval is corroborated by recent studies showing core DMN responds when decisions are guided by working memory (Murphy et al., 2018; 2019), as well as showing stronger responses during autobiographical memory when participants report being more focussed on the task and not distracted by concurrent visual input (Zhang et al., 2021). These findings taken together indicate that core DMN is not task-negative, but supports states in which attention is directed towards internally-maintained information.

The MT subsystem showed stronger activation for autobiographical memory, pictorial semantic judgements, and semantic decisions following scene cues. These findings are consistent with the view that this subsystem is recruited when integrating visual-spatial information, and contributes to scene construction (Andrews-Hanna and Grilli, 2021). The MT subsystem reliably showed activation in bilateral medial temporal regions, consistent with previous studies showing these regions are important for thinking about events that happened in a specific place and time during episodic retrieval (Hassabis & Maguire, 2007; Hassabis, Kumaran, and Maguire, 2007). Moreover, intrinsic connectivity of activated regions within MT revealed stronger coupling with the visual end of the second gradient than other DMN subsystems. The role of MT in visual-spatial tasks might be enhanced by this subsystem’s relative proximity to unimodal and visual ends of the gradients, given that activating voxels were closer to visual cortex than deactivating voxels, in a reversal of the pattern for core DMN. Therefore, although core and MT subsystems are both implicated in episodic, internal states such as autobiographical memory retrieval (Andrews-Hanna, 2012), task activation in these subnetworks has opposite relationships with visual coupling.The FT subsystem showed stronger activation for words than pictures and for abstract than concrete words, yet equivalent responses for reading and autobiographical memory, and for semantic decisions following location and emotion cues, suggesting a role in abstract conceptual processing, irrespective of semantic content or whether tasks are internally- or externally-oriented. Activating voxels in FT were situated between MT and Core on the heteromodal-to-unimodal gradient, in line with this subsystem’s recruitment in both externally-driven abstract semantic tasks and internal tasks such as autobiographical memory. Task-induced increases in activation and deactivation were also stronger for more demanding semantic judgements, suggesting that semantic representations are maintained in a “tuned” state when task demands are higher, and suggesting that deactivation is not task-irrelevant; instead, deactivation could suppress irrelevant features, input modalities and patterns of connectivity (Gouws et al., 2014; Schwartz et al., 2007; Stiernman et al., 2021); or improve the processing efficiency and/or reduce the physiological cost of task responses (Szinte and Knapen, 2020). Left AG within the FT subsystem showed the most reliable activation and deactivation overlap across tasks, indicating the sensitivity of this site to changing task demands: this region may support diverse tasks by flexibly gating its connections, consistent with previous evidence showing it contains ‘echoes’ of many other networks and dynamically modulates its response to suit the context (Braga et al., 2013; Spreng et al., 2013; Vatansever et al., 2015; Dixon et al., 2017, 2018; Wang et al., 2021).

Interestingly, we observed that the presentation of irrelevant information (effects of conflict in Study 1 and irrelevant cues in Study 3) did not modulate the response of the DMN subsystems, while the difficulty of semantic retrieval (effects of association strength in Study 2 and abstractness in Study 3) generated both increases *and* decreases in activation in FT. This corroborates the growing view that DMN deactivation cannot be distilled down to difficulty, but is more likely to reflect selective integration of relevant semantic information (Smallwood et al., 2021), which is needed to identify a specific linking context for weak associations or the precise meaning of an abstract word.

There are of course some important limitations of this research. First, we used a restricted range of tasks that focussed on semantic cognition; future research is needed to establish if the pattern observed here is replicated in other domains, such as episodic memory and social cognition. Given that we found involvement of all three subsystems in semantic cognition, with variation across them reflecting representational and input processing demands, we might expect parallel findings in any domain in which these cognitive dimensions can be manipulated, in line with our observation that MT supports picture and scene-based semantic processing as well as episodic memory, while FT supports both reading and autobiographical memory. Another limitation is that different participants were tested on each task; therefore, we cannot explore the extent to which different patterns of recruitment reflect task effects as opposed to individual differences. Studies probing DMN recruitment within the same subjects would be able to establish if there are spatially-correlated patterns of activation and deactivation related to perceptual decoupling, visuo-spatial memory and abstract cognition across tasks: for example, do patterns in FT linked to verbal versus picture-based semantic retrieval also predict the differences between abstract and concrete words? Despite these limitations, our study provides important constraints on theories of DMN functioning.

In conclusion, we show that DMN subsystems play complementary roles in semantic cognition that are related to their distinctive connectivity patterns, captured by their location within a multi-dimensional space defined by spatially-overlapping cortical gradients. None of these DMN variants was task negative; instead, their recruitment varied according to the need to allocate attention to external inputs in service of a task, and to represent visuospatial and abstract conceptual information.

## Acknowledgements

This work was supported by the European Research Council (Project ID: 771863 – FLEXSEM to EJ).

