## Supplemental figures and tables for "Distinctive and complementary roles of default mode network subsystems in semantic cognition"

### Neuroimaging data acquisition

#### Study 1 Reading/autobiographical memory task

Structural and functional data were acquired using a 3T GE HDx Excite MRI scanner utilizing an eight-channel phased array head coil. Structural MRI acquisition in all participants was based on a T1-weighted 3D fast spoiled gradient echo sequence (repetition time (TR) = 7.8 s, echo time (TE) = minimum full, flip angle = 20°, matrix size = 256 × 256, 176 slices, voxel size = 1.13 × 1.13 × 1 $mm^3$). The task-based activity was recorded using single-shot 2D gradient-echo-planar imaging sequence with TR = 3 s, TE = minimum full, flip angle = 90°, matrix size = 64 × 64, 60 slices, and voxel size = 3 × 3 × 3 $mm^3$.

#### Study 2 word/picture modalities task

Structural and functional data were acquired with a GE 3 T HDx Excite MRI scanner at the York Neuroimaging Centre (YNiC), in a single scanning session. A Magnex, 8 channel, gradient insert head coil with a birdcage, radio frequency coil tuned to 127.4 MHz was used. A gradient-echo EPI sequence was used to collect data from 39 contiguous axial slices (TR 3 s, TE=25 ms, FOV 260 $mm^2$, matrix size=128×128, slice thickness=3.5 mm). The functional data were co-registered onto structural T1-weighted images with a resolution of 1 mm×1 mm×1 mm (TR=8.03, TE=3.07 ms, FOV 290 mm×290 mm×176 mm, matrix size 256×256×176, slice thickness=1.13 mm×1.13 mm×1 mm). Functional data were additionally co-registered to T1 weighted FLAIR images (5.6 mm×5.6 mm×3.5 mm), taken in the same plane as the EPI slices with interleaved slice acquisition.

#### Study 3 abstract/concrete words task

Images were acquired on a 3T Philips Achieva scanner using an 8 element SENSE head coil with a sense factor of 2.5. A spin-echo imaging sequence, combined with a post-acquisition distortion-correction, was employed to improve signal quality in the vATL (Embleton et al., 2010). The spin-echo EPI sequence included 31 slices covering the whole brain with echo time (TE) = 70 msec, time to repetition (TR) = 3200 msec, flip angle = 90°, 96 × 96 matrix, reconstructed in-plane resolution 2.5 × 2.5 mm, slice thickness 4.0 mm. Following the standard method for distortion-corrected spin-echo fMRI (Embleton et al., 2010), the images were acquired with a single direction k space traversal and a left-right phase encoding direction. In between the two functional runs, a brief "pre-scan" was acquired, consisting of 10 volumes of dual direction k space traversal SE EPI scans. These scans were used in the distortion correction procedure. In addition, a high resolution T1-weighted 3D turbo field echo inversion recovery image was acquired (TR = 8400 msec, TE = 3.9 msec, flip angle 8°, 256 × 205 matrix reconstructed to 256 × 256, reconstructed resolution .938 × .938 mm, and slice thickness of 0.9 mm, SENSE factor = 2.5) with 160 slices covering the whole brain. This image was used for spatial normalisation.

#### Study 4 emotional/spatial cues task

Structural and functional data were acquired with a GE 3 T HDx Excite MRI scanner. Structural MRI data acquisition in all participants was based on a T1-weighted 3D fast spoiled gradient echo sequence (TR = 7.8 ms, TE = minimum full, flip-angle = 20°, matrix size = 256 × 256, 176 slices, voxel size = 1.13 × 1.13 × 1 mm). A gradient-echo EPI sequence was used to collect functional data from 60 interleaved bottom-up axial slices aligned with the temporal lobe (TR = 3s, TE = 18.9 ms, FOV = 192 × 192 × 180 mm, matrix size = 64 × 64, slice thickness = 3 mm, slice-gap = 3 mm, voxel size = 3 × 3x3 mm3, flip-angle = 90°). An intermediary FLAIR scan with the same orientation as the functional scans was collected to improve the co-registration between subject-specific structural and functional scans.

#### Study 5 Resting-state task

Structural and functional MRI data were acquired on a 3T GE HDx Excite MRI scanner, equipped with an eight-channel phased array head coil at the York Neuroimaging Centre, University of York. For each participant, structural MRI was acquired based on a sagittal isotropic 3D fast spoiled gradient-recalled echo T1-weighted structural scan (TR = 7.8 ms, TE = minimum full, flip angle = 20°, matrix size = 256 × 256, 176 slices, voxel size = 1.13 mm × 1.13 mm × 1 mm). The 9-

minute resting-state fMRI data were acquired using a gradient single-shot 10 echo-planar imaging sequence (TE = minimum full, flip angle = 90°, matrix = 64 × 64, FOV = 192mm × 192 mm, voxel size = 3 mm × 3 mm × 3 mm, TR = 3000 ms, 60 slices with no gap).

#### Pre-processing and individual-level analysis of fMRI data

##### Study 1 Reading/autobiographical memory task

All functional and structural data were preprocessed using a standard pipeline and analysed via the FMRIB Software Library (FSL version 6.0, www.fmrib.ox.ac.uk/fsl). Individual T1-weighted structural brain images were extracted using FSL’s Brain Extraction Tool (BET). Structural images were linearly registered to the MNI152 template using FMRIB's Linear Image Registration Tool (FLIRT). The first three volumes of each functional scan were removed in order to minimise the effects of magnetic saturation. The functional neuroimaging data were analysed using FSL’s FMRI Expert Analysis Tool (FEAT). We applied motion correction using MCFLIRT (37), slice-timing correction using Fourier space time-series phase-shifting (interleaved), spatial smoothing using a Gaussian kernel of FWHM 6 mm, and high-pass temporal filtering (sigma = 100 s) to remove temporal signal drift. In addition, motion scrubbing (using the fsl_motion_outliers tool) was applied to exclude volumes that exceeded a framewise displacement threshold of 0.9.

The pre-processed time-series data were modelled using a general linear model, using FMRIB’s Improved Linear Model (FILM) correcting for local autocorrelation. Nine Explanatory Variables (EV) of interest and nine of no interest were modelled using a double-Gaussian hemodynamic response gamma function. The nine EVs of interest were: Reading (1) without and (2) with conflict from memory recall, Autobiographical memory retrieval (3) with and (4) without conflict from semantic input, (5) Letter String Baseline, (6-9) Task Focus effect for each of the four experimental conditions as a parametric regressor. Our EVs of no interest were: (10) Memory cue words and (11) Letter strings before the presentation of task instructions, Task instructions for Reading (12) without and (13) with conflict (this separation of the reading task instruction was based on the consideration that some recall or task preparation was likely to be occurring due to the presentation of autobiographical memory cues), plus task instructions for (14) Memory Recall and (15) Letter String baseline conditions. Other EVs of no interest were: (16) Fixation (the inter-stimulus fixations between the sentences or letter strings and the ratings questions), (17) Responses to catch trials (which included all time points with responses across conditions), and (18) Rating decision periods (including all the ratings across experimental conditions). EVs for each condition commenced at the onset of the first word of the sentence or the first letter string, with EV duration set as the presentation time (9s). The parametric EVs for the effect of Task Focus during the target had the same onset time and duration as the EVs corresponding to the four experimental trials, but in addition included the demeaned Task Focus ratings value as a weight. The fixation period between the trials provided the implicit baseline. We examined the main effects of Task, and Conflict for both the main experimental conditions compared with the implicit baseline, which allowed us to identify the activation and deactivation in each task.

#### Study 2 word/picture modalities task

We used an event-related design for all of the analyses (i.e., to examine the effects of both difficulty and task), even though the various tasks were presented in mini-blocks. Only accurate responses were used in the analysis. All first-level and higher-level analyses were run using FMRI Expert Analysis Tool (FEAT) Version 5.98, in FMRIB's Software Library (FSL), www.fmrib.ox.ac.uk/fsl). Prior to inferential statistical analysis the following pre-processing was applied: Individual brain extraction (BET) to remove non-brain material from images for co-registration of the functional data, MCFLIRT motion correction (using fMRIB's Linear Registration Tool; Jenkinson et al., 2002), slice timing correction using Fourier-space time-series phase shifting (Sinc interpolation with a Hanning-windowing kernel), FWHM 6.0 mm spatial smoothing (Gaussian Kernel), high-pass temporal filtering (Gaussian-weighted least-squares straight line fitting, with sigma=100 s). We used FILM nonparametric estimation of time series autocorrelation (FILM; FMRIB's Improved Linear Model) to fit the model to the data, on all lower-level analyses. FSL's canonical gamma HRF along with a temporal derivative was used to model the HRF response. The first two volumes were removed to allow for T1 saturation effects. To analyse the data at the group level, we entered lower level FEAT directories into a higher level FMRIB'S Local Analysis of Mixed Effects (FLAME) Bayesian mixed effects analysis (Beckmann et al., 2003, Woolrich, 2008, Woolrich et al., 2004). Z (Gaussianised T/F) statistic images were thresholded using clusters determined by Z>2.3 and a (corrected) cluster significance threshold of p<.05 (Worsley, 2001). Names of brain areas reported are labelled according to the Harvard-Oxford Cortical Structural Atlas, Talairach Deamon and the Juelich Histological Atlas built into the FSLView software library.

Each task and condition was modelled separately using event based explanatory variables (EV) which were convolved to the haemodynamic response function (gamma function). We used a variable-epoch model as recommended by Grinband et al. (2008) to capture effects of time-on-task within each EV: the haemodynamic response function was aligned to the beginning of each correct trial and lasted for the duration of the event. Incorrect/removed trials were modelled as a separate EV, therefore, any data not modelled was included as rest. Several contrasts were run (11 in total): A contrast against rest/baseline was conducted for each of the six conditions (non-semantic easy, non-semantic hard, semantic verbal easy, semantic verbal hard, semantic picture easy, semantic picture hard), the hard version of each judgement type was contrasted against the corresponding easy version (non-semantic hard–non-semantic easy, semantic verbal hard–semantic verbal easy, etc.), and two contrasts examining modality/task were included (semantic verbal-rhyme; semantic picture–semantic verbal).

#### Study 3 abstract/concrete words task

Analysis was carried out using SPM8. The motion and distortion-corrected images for each participant were first co-registered to their T1 structural scan. Spatial normalisation of the T1 scans into MNI space was computed using DARTEL (Ashburner, 2007) and the resulting transformation applied to the functional images, which were resampled to 2 × 2 × 2 mm voxel size and smoothed with an 8 mm FWHM Gaussian kernel. At this point, temporal signal-to-noise (TSNR) maps were generated for each participant by dividing the mean signal in each voxel by its standard deviation (Murphy, Bodurka, & Bandettini, 2007). The mean TSNR map across all participants is shown in Fig. 1. TSNR exceeded 80 in ventral temporal regions. Unlike gradient-echo fMRI, which shows a pronounced drop in TSNR in ventral temporal regions relative to the rest of the brain, TSNR in the ventral temporal lobes was comparable to that in frontal and superior temporal regions. The data were treated with a high-pass filter with a cut-off of 190 sec and analysed using a general linear model. At the first level, each of the five stimulus conditions was modelled with a separate regressor (concrete-context, concrete-irrelevant, abstract-context, abstract-irrelevant and number baseline). Blocks were modelled with a boxcar function convolved with the canonical haemodynamic response function. Motion parameters were entered into the model as covariates of no interest. Parameter estimates were subjected to several analyses, each targeted at a specific hypothesis.

#### Study 4 emotional/spatial cues task

FMRI data processing was carried out using FEAT (FMRI Expert Analysis Tool) Version 6.0, part of FSL (FMRIB's Software Library, www.fmrib.ox.ac.uk/fsl). Registration of the high resolution structural to standard space (Montreal Neurological Institute – MNI) was carried out using FLIRT (Jenkinson et al., 2002; Jenkinson and Smith, 2001). Pre-processing of the functional image included motion correction using MCFLIRT (Jenkinson et al., 2002), slice-timing correction using Fourier-space time-series phase-shifting (interleaved), non-brain removal using BET (Smith, 2002), spatial smoothing using a Gaussian kernel of FWHM 5 mm, grand-mean intensity normalisation of the entire 4D dataset by a single multiplicative factor, and high-pass temporal filtering (Gaussian-weighted least-squares straight line fitting, with sigma = 50.0s).

#### Study 5 Resting-state task

fMRI data was pre-processed using SPM12 (http://www.fil.ion.ucl.ac.uk/spm) and CONN (v.18b) (https://www.nitrc.org/projects/conn) (Whitfield-Gabrieli & Nieto-Castanon, 2012) implemented in Matlab (R2018a) (https://uk.mathworks.com/products/matlab). Pre-processing steps followed CONN's default pipeline and included motion estimation and correction by volume realignment using a six-parameter rigid body transformation, slice-time correction, and simultaneous grey matter (GM), white matter (WM) and cerebrospinal fluid (CSF) segmentation and normalisation to MNI152 stereotactic space (2 mm isotropic) of both functional and structural data. Following pre-processing, the following potential confounds were statistically controlled for: 6 motion parameters calculated at the previous step and their 1st and 2nd order derivatives, volumes with excessive movement (motion greater than 0.5 mm and global signal changes larger than z = 3), linear drifts, and five principal components of the signal from WM and CSF (CompCor approach; (Behzadi, Restom, Liau, & Liu, 2007). Finally, data were band-pass filtered between 0.01 and 0.1 Hz. No global signal regression was performed (Vos de Wael, Hyder, & Thompson, 2017).

#### Supplementary figures and tables

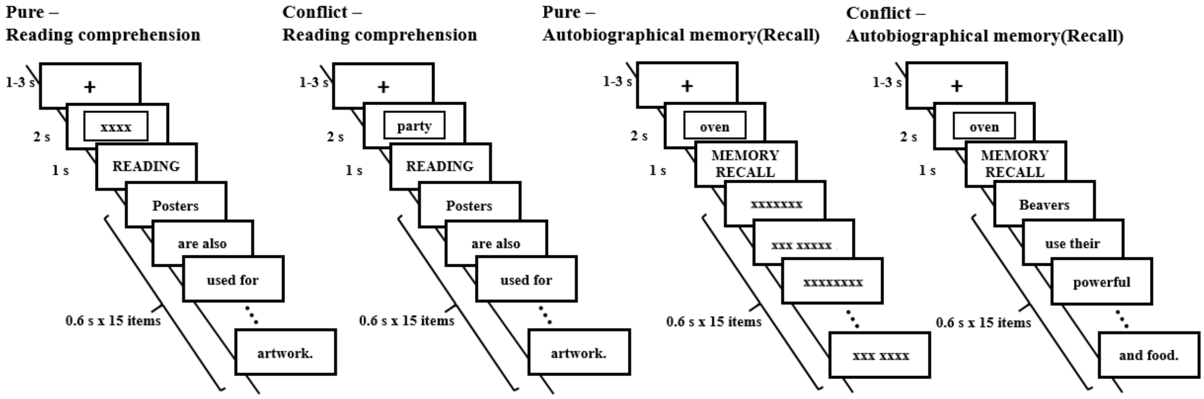


**Figure S1** Task illustration of normal and conflict conditions for the reading comprehension and autobiographical memory tasks (Study 1).

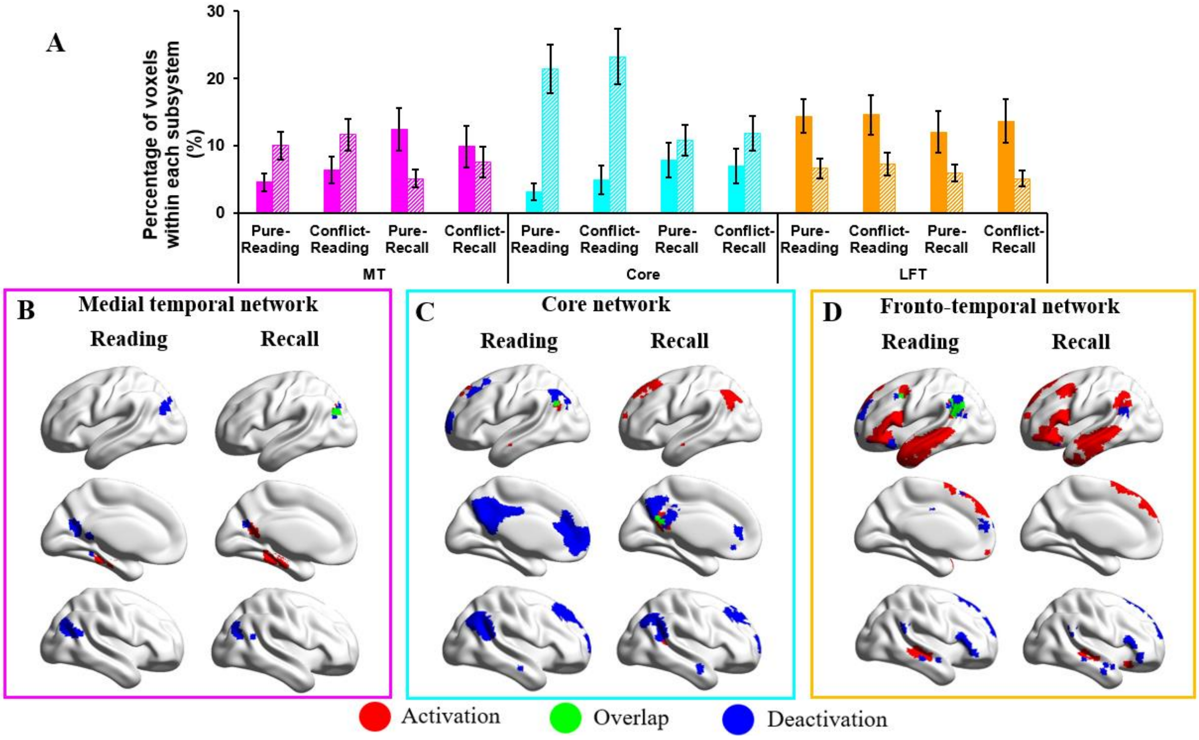


**Figure S2 Panel A** shows the percentage of activating or deactivating voxels in conflict/no-conflict reading or autobiographical memory (recall) tasks (Study 1). The three colours represent the percentage of voxels extracted from three subsystems defined by Yeo et al. (2011) in their 17-network parcellation (pink = medial temporal regions, cyan = core regions, orange = fronto-temporal regions). Solid bars represent the percentages of activating voxels, and grid bars represent the percentage of deactivating voxels. Error bars represent one standard error. **Panels B-D** show the regions of activating voxels (in red colour), deactivating voxels (in blue colour), and the overlapping regions of both activating and deactivating voxels (i.e., regions showed activation in several participants, and deactivation in another group of participants, in green colour) in reading or recall conditions in MT, core and FT subsystems, respectively.

**Table S1** Results of repeated-measured ANOVAs for percentage of activation and deactivation voxels in conflict/no-conflict reading or recall conditions (Study 1)

| | | **Percentage of activation voxels** | | | **Percentage of deactivation voxels** | | |
|---|---|---|---|---|---|---|---|
| | | F | p | $\eta^2$ | F | p | $\eta^2$ |
| **Main effect** | Task (Reading/Recall) | F(1,28) = 5.36 | **.028** | .16 | F(1,28) = 28.13 | **< .001** | .50 |
| | Task Conflict (Pure/Conflict) | F(1,28) = .26 | .62 | .01 | F(1,28) = 1.99 | .17 | .07 |
| | DMN | F(2,56) = 22.14 | **< .001** | .44 | F(2,56) = 22.01 | **< .001** | .44 |
| **Two-way interaction** | Task × Task Conflict | F(1,28) = 1.65 | .21 | .06 | F(1,28) = .09 | .76 | .003 |
| | Task × DMN | F(2,56) = 22.18 | **< .001** | .44 | F(2,56) = 21.94 | **< .001** | .44 |
| | Task Conflict × DMN | F(2,56) = 1.56 | .22 | .05 | F(2,56) = 2.66 | .08 | .09 |
| **Three-way interaction** | Task × Task Conflict × DMN | F(2,56) = 9.41 | **< .001** | .25 | F(2,56) = 1.04 | .36 | .04 |

**Table S2** Results of repeated-measured ANOVAs for percentage of activation and deactivation voxels in conflict/no-conflict reading or recall conditions in each DMN subsystem (Study 1)

| **Percentage of activation voxels** | | | | | | | | | | |
|---|---|---|---|---|---|---|---|---|---|---|
| | | **Medial temporal subsystem** | | | **Core subsystem** | | | **Fronto-temporal subsystem** | | |
| | | F | p | $\eta^2$ | F | p | $\eta^2$ | F | p | $\eta^2$ |
| **Main effect** | Task (Reading/Recall) | F(1,28) = 11.87 | **.002** | .30 | F(1,28) = 10.24 | **.003** | .27 | F(1,28) = 3.24 | .08 | .10 |
| | Task Conflict (Pure/Conflict) | F(1,28) = 0.14 | .71 | .005 | F(1,28) = .62 | .44 | .02 | F(1,28) = 1.34 | .26 | .05 |
| **Two-way interaction** | Task × Task Conflict | F(1,28) = 6.38 | **.017** | .19 | F(1,28) = 2.07 | .16 | .07 | F(1,28) = 1.10 | .30 | .04 |

| **Percentage of deactivation voxels** | | | | | | | | | | |
|---|---|---|---|---|---|---|---|---|---|---|
| | | **Medial temporal subsystem** | | | **Core subsystem** | | | **Fronto-temporal subsystem** | | |
| | | F | p | $\eta^2$ | F | p | $\eta^2$ | F | p | $\eta^2$ |
| **Main effect** | Task (Reading/Recall) | F(1,28) = 17.49 | **≤ .001** | .39 | F(1,28) = 30.98 | **≤ .001** | .53 | F(1,28) = 5.36 | **.044** | .14 |
| | Task Conflict (Pure/Conflict) | F(1,28) = 3.82 | .061 | .12 | F(1,28) = 1.37 | .25 | .05 | F(1,28) = .26 | .91 | .00 |
| **Two-way interaction** | Task × Task Conflict | F(1,28) = .23 | .64 | .008 | F(1,28) = .095 | .76 | .003 | F(1,28) = 1.65 | .14 | .08 |

**Table S3** Results of repeated-measured ANOVAs for percentage of activation and deactivation voxels in easy/hard verbal or picture conditions (Study 2)

| | | Percentage of activation voxels | | | Percentage of deactivation voxels | | |
|---|---|---|---|---|---|---|---|
| | | F | p | $\eta^2$ | F | p | $\eta^2$ |
| **Main effect** | Modality (verbal/picture) | F(1,21) = 2.33 | .142 | .10 | F(1,21) = 4.70 | **.042** | .18 |
| | Difficulty (easy/hard) | F(1,21) = 15.50 | **< .001** | .43 | F(1,21) = 25.73 | **< .001** | .55 |
| | DMN | F(2,42) = 33.31 | **< .001** | .61 | F(2,42) = 122.75 | **< .001** | .85 |
| **Two-way interaction** | Modality × Difficulty | F(1,21) = 1.67 | .21 | .07 | F(1,21) = .63 | .44 | .03 |
| | Modality × DMN | F(2,42) = 73.32 | **< .001** | .78 | F(2,42) = 33.14 | **< .001** | .61 |
| | Difficulty × DMN | F(2,42) = 10.63 | **< .001** | .34 | F(2,42) = 9.18 | **< .001** | .30 |
| **Three-way interaction** | Modality × Difficulty × DMN | F(2,42) = .73 | .49 | .03 | F(2,42) = 1.96 | .15 | .09 |

**Table S4** Results of repeated-measured ANOVAs for percentage of activation and deactivation voxels in easy/hard verbal or picture conditions in each DMN subsystem (Study 2)

**Percentage of activation voxels**

| | | **Medial temporal subsystem** | | | **Core subsystem** | | | **Fronto-temporal subsystem** | | |
|---|---|---|---|---|---|---|---|---|---|---|
| | | F | p | $\eta^2$ | F | p | $\eta^2$ | F | p | $\eta^2$ |
| **Main effect** | Modality (verbal/picture) | F(1,21) = 38.33 | **<u>< .001</u>** | .65 | F(1,21) = 1.09 | .31 | .05 | F(1,21) = 119.29 | **<u>< .001</u>** | .85 |
| | Difficulty (easy/hard) | F(1,21) = 3.33 | .08 | .14 | F(1,21) = 3.91 | .061 | .16 | F(1,21) = 35.69 | **<u>< .001</u>** | .63 |
| **Two-way interaction** | Modality × Difficulty | F(1,21) = 2.26 | .15 | .10 | F(1,21) = 3.06 | .095 | .13 | F(1,21) = .10 | .76 | .005 |

**Percentage of deactivation voxels**

| | | **Medial temporal subsystem** | | | **Core subsystem** | | | **Fronto-temporal subsystem** | | |
|---|---|---|---|---|---|---|---|---|---|---|
| | | F | p | $\eta^2$ | F | p | $\eta^2$ | F | p | $\eta^2$ |
| **Main effect** | Modality (verbal/picture) | F(1,21) = 6.43 | **<u>.019</u>** | .23 | F(1,21) = 8.26 | **<u>.009</u>** | .28 | F(1,21) = 31.33 | **<u>< .001</u>** | .60 |
| | Difficulty (easy/hard) | F(1,21) = 28.49 | **<u>< .001</u>** | .58 | F(1,21) = 20.30 | **<u>< .001</u>** | .49 | F(1,21) = 5.37 | **<u>.031</u>** | .20 |
| **Two-way interaction** | Modality × Difficulty | F(1,21) = 1.90 | .18 | .08 | F(1,21) = .37 | .55 | .02 | F(1,21) = 0.00 | .99 | 0.00 |

**Table S5** Examples of the cueing sentences and the semantic judgement task (Study 3)

| Condition | Cue | Judgement |
|---|---|---|
| **Contextual cue – abstract words** | The road is closed.<br>We must look for an alternative. | alternative<br>substitute ambition discretion |
| **Contextual cue – concrete words** | I ordered the roast dinner.<br>It was served with asparagus. | asparagus<br>broccoli christmas furniture |
| **Irrelevant cue – abstract words** | He's very ignorant.<br>He never takes a hint. | rate<br>worth speed excuse |
| **Irrelevant cue – concrete words** | We had sandwiches for lunch.<br>They contained cucumber. | villain<br>herring crook aluminium |

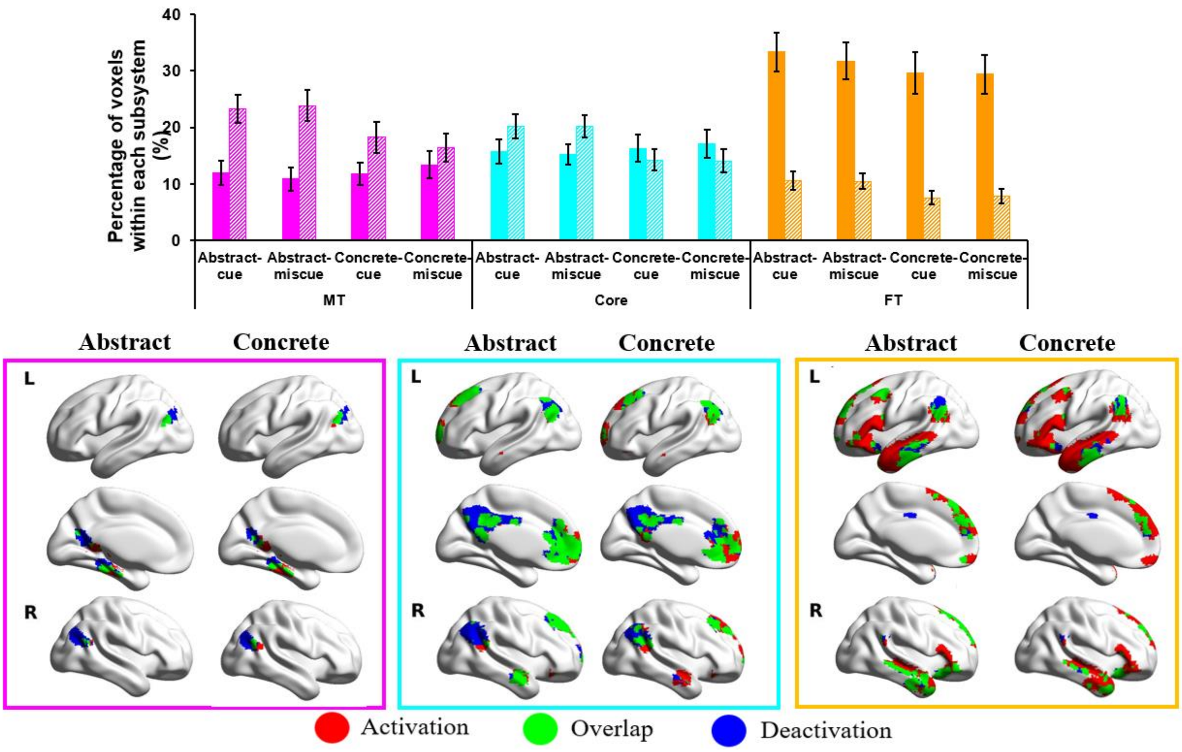


**Figure S3 Panel A** shows the percentage of activating or deactivating voxels in abstract or concrete conditions with contextual or irrelevant cues (Study 3; cue = contextual cue, miscue = irrelevant cue). The three colours represent the percentage of voxels extracted from three subsystems defined by Yeo et al. (2011) in their 17-network parcellation (pink = medial temporal regions, cyan = core regions, orange = fronto-temporal regions). Solid bars represent the percentages of activating voxels, and grid bars represent the percentage of deactivating voxels. Error bars represent one standard error. **Panels B-D** show the regions of activating voxels (in red colour), deactivating voxels (in blue colour), and the overlapping regions of both activating and deactivating voxels (i.e., regions showed activation in several participants, and deactivation in another group of participants, in green colour) in abstract or concrete conditions in MT, core, and FT subsystems, respectively.

**Table S6** Results of repeated-measured ANOVAs for percentage of activation and deactivation voxels in relevant-cue/irrelevant-cue abstract or concrete conditions (Study 3)

| | | **Percentage of activation voxels** | | | **Percentage of deactivation voxels** | | |
|---|---|---|---|---|---|---|---|
| | | F | p | $\eta^2$ | F | p | $\eta^2$ |
| **Main effect** | Concreteness (Abstract/concrete) | F(1,18) = .10 | .75 | .006 | F(1,18) = 59.90 | **< .001** | .77 |
| | Cue (relevant/irrelevant cue) | F(1,18) = .03 | .86 | .002 | F(1,18) = .15 | .70 | .008 |
| | DMN | F(2,36) = 30.02 | **< .001** | .63 | F(2,36) = 20.73 | **< .001** | .54 |
| **Two-way interaction** | Concreteness × Cue | F(1,18) = 2.28 | .15 | .11 | F(1,18) = .33 | .58 | .02 |
| | Concreteness × DMN | F(2,36) = 13.45 | **< .001** | .43 | F(2,36) = 5.86 | **.018** | .25 |
| | Cue × DMN | F(2,36) = 1.10 | .34 | .06 | F(2,36) = .34 | .71 | .02 |
| **Three-way interaction** | Concreteness × Cue × DMN | F(2,36) = .48 | .57 | .03 | F(2,36) = .84 | .41 | .04 |

**Table S7** Results of repeated-measured ANOVAs for percentage of activation and deactivation voxels in relevant-cue/irrelevant-cue abstract or concrete conditions in each DMN subsystem (Study 3)

**Percentage of activation voxels**

| | | **Medial temporal subsystem** | | | **Core subsystem** | | | **Fronto-temporal subsystem** | | |
|---|---|---|---|---|---|---|---|---|---|---|
| | | F | p | $\eta^2$ | F | p | $\eta^2$ | F | p | $\eta^2$ |
| **Main effect** | Concreteness (Abstract/concrete) | F(1,18) = 3.17 | .09 | .15 | F(1,18) = 1.49 | .24 | .08 | F(1,18) = 14.47 | **<u>< .001</u>** | .45 |
| | Cue (relevant/irrelevant cue) | F(1,18) = .12 | .73 | .007 | F(1,18) = .03 | .87 | .002 | F(1,18) = .75 | .40 | .04 |
| **Two-way interaction** | Concreteness × Cue | F(1,18) = 3.92 | .063 | .18 | F(1,18) = .84 | .37 | .04 | F(1,18) = .63 | .44 | .03 |

**Percentage of deactivation voxels**

| | | **Medial temporal subsystem** | | | **Core subsystem** | | | **Fronto-temporal subsystem** | | |
|---|---|---|---|---|---|---|---|---|---|---|
| | | F | p | $\eta^2$ | F | p | $\eta^2$ | F | p | $\eta^2$ |
| **Main effect** | Concreteness (Abstract/concrete) | F(1,18) = 21.90 | **<u>< .001</u>** | .55 | F(1,18) = 84.00 | **<u>< .001</u>** | .82 | F(1,18) = 30.73 | **<u>< .001</u>** | .63 |
| | Cue (relevant/irrelevant cue) | F(1,18) = .34 | .57 | .02 | F(1,18) = .05 | .83 | .003 | F(1,18) = .02 | .88 | .001 |
| **Two-way interaction** | Concreteness × Cue | F(1,18) = .97 | .34 | .05 | F(1,18) = .004 | .95 | .000 | F(1,18) = .06 | .80 | .004 |

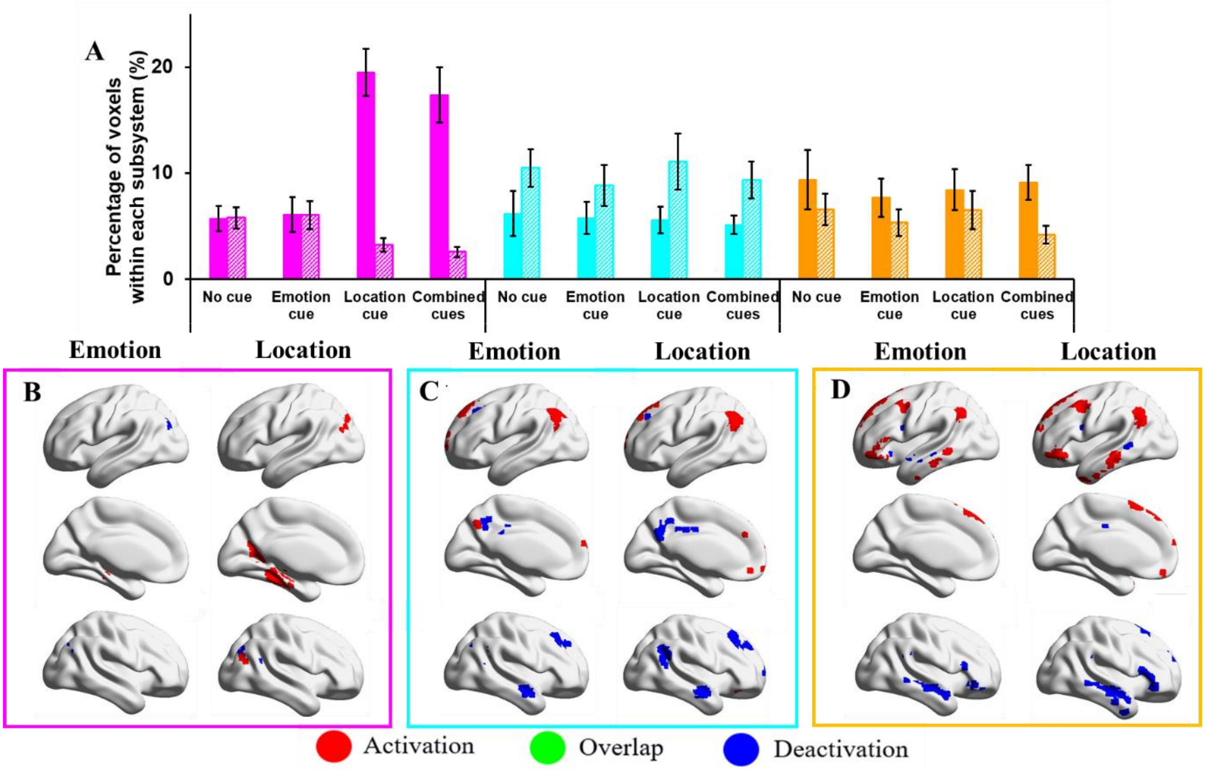


**Figure S4 Panel A** shows the percentage of activating or deactivating voxels in conditions with 1) no cue; 2) emotional cue; 3) spatial cue; or 4) combined cues (Study 4). The three colours represent the percentage of voxels extracted from three subsystems defined by Yeo et al. (2011) in their 17-network parcellation (pink = medial temporal regions, cyan = core regions, orange = fronto-temporal regions). Solid bars represent the percentages of activating voxels, and grid bars represent the percentage of deactivating voxels. Error bars represent one standard error. **Panels B-D** show the regions of activating voxels (in red colour), deactivating voxels (in blue colour), and the overlapping regions of both activating and deactivating voxels (i.e., regions showed activation in several participants, and deactivation in another group of participants, in green colour) in emotional-cue or spatial-cue conditions in MT, core, and FT subsystems, respectively.

**Table S8** Results of repeated-measured ANOVAs for percentage of activation and deactivation voxels in no cue, emotional/spatial cue, or combined cues conditions in each DMN subsystem (Study 4)

| | | **Percentage of activation voxels** | | | **Percentage of deactivation voxels** | | |
|---|---|---|---|---|---|---|---|
| | | F | p | $\eta^2$ | F | p | $\eta^2$ |
| **Main effect** | Cue (no cue, emotional/spatial cue, or combined cues) | 3.62 | **.017** | .13 | 1.25 | .30 | .05 |
| | DMN | 10.26 | **.001** | .29 | 12.52 | **< .001** | .33 |
| **Two-way interaction** | Cue × DMN | 20.32 | **< .001** | .45 | 2.21 | .085 | .08 |

**Table S9** Results of paired t-tests (Bonferroni corrected for the number of comparisons) for percentage of activation and deactivation voxels between no cue, emotional (E)/spatial (S) cue, or combined (C) cues within MT subsystem for activation and deactivation respectively (Study 4).

| | | **Activation within each subsystem** | | | **Deactivation within each subsystem** | | |
|---|---|---|---|---|---|---|---|
| | | *df* | *t* | *p* | *df* | *t* | *p* |
| **MT** | 0 cue vs. E cue | 25 | - .22 | *.83* | 25 | -.21 | *.84* |
| | 0 cue vs. S cue | 25 | -5.43 | **< .001** | 25 | 2.54 | .072 |
| | E cue vs. C cues | 25 | -4.34 | **< .001** | 25 | 3.14 | **.016** |
| | S cue vs. C cues | 25 | 1.07 | .30 | 25 | .92 | .37 |

**Table S10** Results of repeated-measured ANOVAs for percentage of activation and deactivation voxels thresholded at z=2.3/2.6 in reading or recall conditions (Study 1)

| **Threshold z=2.3** | **Activation ANOVA** | | | | **Deactivation ANOVA** | | | |
|---|---|---|---|---|---|---|---|---|
| | *df* | *F* | *p* | $\eta^2$ | *df* | *F* | *p* | $\eta^2$ |
| **Task (Reading/Recall)** | (1, 28) | 3.66 | .066 | .12 | (1, 28) | 24.15 | **< .001** | .46 |
| **DMN** | (2, 56) | 17.90 | **< .001** | .39 | (2, 56) | 18.47 | **< .001** | .40 |
| **Task × DMN** | (2, 56) | 20.71 | **< .001** | .43 | (2, 56) | 20.00 | **< .001** | .42 |
| **Threshold z=2.6** | **Activation ANOVA** | | | | **Deactivation ANOVA** | | | |
| | *df* | *F* | *p* | $\eta^2$ | *df* | *F* | *p* | $\eta^2$ |
| **Task (Reading/Recall)** | (1, 28) | 2.42 | .131 | .08 | (1, 28) | 20.42 | **< .001** | .42 |
| **DMN** | (2, 56) | 14.34 | **< .001** | .34 | (2, 56) | 16.04 | **< .001** | .36 |
| **Task × DMN** | (2, 56) | 21.34 | **< .001** | .43 | (2, 56) | 17.53 | **< .001** | .39 |

**Table S11** Results of paired t-tests (Bonferroni corrected for the number of comparisons) between reading and recall conditions within each subsystem for activation and deactivation respectively (Study 1).

| **Threshold z=2.3** | **Activation within each subsystem** | | | **Deactivation within each subsystem** | | |
|---|---|---|---|---|---|---|
| | *df* | *t* | *p* | *df* | *t* | *p* |
| **MT** | 28 | 3.04 | **.015** | 28 | 3.76 | **< .001** |
| **Core** | 28 | 2.83 | **.027** | 28 | 5.17 | **< .001** |
| **FT** | 28 | 1.83 | .234 | 28 | 2.36 | .078 |
| **Threshold z=2.6** | **Activation within each subsystem** | | | **Deactivation within each subsystem** | | |
| | *df* | *t* | *p* | *df* | *t* | *p* |
| **MT** | 28 | -2.80 | **.027** | 28 | 3.42 | **.006** |
| **Core** | 28 | -2.59 | **.045** | 28 | 4.76 | **< .001** |
| **FT** | 28 | 1.90 | .204 | 28 | 2.38 | .072 |

**Table S12** Results of repeated-measured ANOVAs for percentage of activation and deactivation voxels thresholded at z=2.3/2.6 in easy/hard verbal or picture matching tasks (Study 2).

| **Threshold z=2.3** | **Percentage of activation voxels** | | | | **Percentage of deactivation voxels** | | | |
|---|---|---|---|---|---|---|---|---|
| | *df* | *F* | *p* | $\eta^2$ | *df* | *F* | *p* | $\eta^2$ |
| **Modality (verbal/picture)** | (1,21) | 1.95 | .18 | .09 | (1,21) | 6.09 | **.022** | .23 |
| **Difficulty (easy/hard)** | (1,21) | 18.97 | **< .001** | .48 | (1,21) | 36.87 | **< .001** | .64 |
| **DMN** | (2,42) | 28.21 | **< .001** | .57 | (2,42) | 100.39 | **< .001** | .83 |
| **Modality × Difficulty** | (1,21) | 1.96 | .18 | .09 | (1,21) | .27 | .61 | .01 |
| **Modality × DMN** | (2,42) | 62.77 | **< .001** | .75 | (1,21) | 26.79 | **< .001** | .56 |
| **Difficulty × DMN** | (2,42) | 9.96 | **< .001** | .32 | (2,42) | 12.76 | **< .001** | .38 |
| **Modality × Difficulty × DMN** | (2,42) | 2.22 | .12 | .10 | (2,42) | 2.15 | .13 | .09 |
| **Threshold z=2.6** | **Percentage of activation voxels** | | | | **Percentage of deactivation voxels** | | | |
| | *df* | *F* | *p* | $\eta^2$ | *df* | *F* | *p* | $\eta^2$ |
| **Modality (verbal/picture)** | (1,21) | 1.75 | .20 | .08 | (1,21) | 7.03 | **.015** | .25 |
| **Difficulty (easy/hard)** | (1,21) | 20.63 | **< .001** | .50 | (1,21) | 46.85 | **< .001** | .69 |
| **DMN** | (2,42) | 24.27 | **< .001** | .54 | (2,42) | 84.23 | **< .001** | .80 |
| **Modality × Difficulty** | (1,21) | 1.90 | .18 | .08 | (1,21) | .06 | .82 | .00 |
| **Modality × DMN** | (2,42) | 55.71 | **< .001** | .73 | (1,21) | 21.50 | **< .001** | .51 |
| **Difficulty × DMN** | (2,42) | 9.97 | **< .001** | .32 | (2,42) | 18.29 | **< .001** | .47 |
| **Modality × Difficulty × DMN** | (2,42) | 3.54 | **.038** | .14 | (2,42) | 1.97 | .15 | .09 |

**Table S13** Results of two-way repeated-measured ANOVAs for easy/hard verbal or picture conditions, within each subsystem and of activation and deactivation voxels thresholded at z=2.3/2.6 respectively (Study 2).

| | | | | | | | | | | | | |
|---|---|---|---|---|---|---|---|---|---|---|---|---|
| | | | | | **Percentage of activation voxels** | | | | | | | |
| **Threshold z=2.3** | **Medial temporal subsystem** | | | | **Core subsystem** | | | | **Fronto-temporal subsystem** | | | |
| | *df* | *F* | *p* | $\eta^2$ | *df* | *F* | *p* | $\eta^2$ | *df* | *F* | *p* | $\eta^2$ |
| **Modality (verbal/picture)** | (1,21) | 32.62 | **< .001** | .61 | (1,21) | .91 | .352 | .04 | (1,21) | 101.59 | **< .001** | .83 |
| **Difficulty (easy/hard)** | (1,21) | 4.86 | **.039** | .19 | (1,21) | 4.48 | **.046** | .18 | (1,21) | 36.94 | **< .001** | .64 |
| **Modality × Difficulty** | (1,21) | 4.00 | .059 | .16 | (1,21) | 3.15 | .091 | .13 | (1,21) | .002 | .97 | .00 |
| | | | | | **Percentage of deactivation voxels** | | | | | | | |
| **Threshold z=2.3** | **Medial temporal subsystem** | | | | **Core subsystem** | | | | **Fronto-temporal subsystem** | | | |
| | df | F | p | $\eta^2$ | df | F | p | $\eta^2$ | df | F | p | $\eta^2$ |
| **Modality (verbal/picture)** | (1,21) | 4.06 | .057 | .16 | (1,21) | 9.73 | **.005** | .32 | (1,21) | 29.82 | **< .001** | .59 |
| **Difficulty (easy/hard)** | (1,21) | 35.14 | **< .001** | .63 | (1,21) | 31.08 | **< .001** | .60 | (1,21) | 8.22 | **.009** | .28 |
| **Modality × Difficulty** | (1,21) | 1.50 | .23 | .07 | (1,21) | .14 | .71 | .01 | (1,21) | .15 | .70 | .01 |
| | | | | | **Percentage of activation voxels** | | | | | | | |
| **Threshold z=2.6** | **Medial temporal subsystem** | | | | **Core subsystem** | | | | **Fronto-temporal subsystem** | | | |
| | *df* | *F* | *p* | $\eta^2$ | *df* | *F* | *p* | $\eta^2$ | *df* | *F* | *p* | $\eta^2$ |
| **Modality (verbal/picture)** | (1,21) | 28.75 | **< .001** | .58 | .(1,21) | .81 | .379 | .04 | (1,21) | 89.33 | **< .001** | .81 |
| **Difficulty (easy/hard)** | (1,21) | 5.71 | **.026** | .21 | (1,21) | 4.37 | **.049** | .17 | (1,21) | 36.59 | **< .001** | .64 |
| **Modality × Difficulty** | (1,21) | 4.83 | **.039** | .19 | (1,21) | 3.12 | .092 | .13 | (1,21) | .15 | .70 | .01 |
| | | | | | **Percentage of deactivation voxels** | | | | | | | |
| **Threshold z=2.6** | **Medial temporal subsystem** | | | | **Core subsystem** | | | | **Fronto-temporal subsystem** | | | |
| | df | F | p | $\eta^2$ | df | F | p | $\eta^2$ | df | F | p | $\eta^2$ |
| **Modality (verbal/picture)** | (1,21) | 2.35 | .14 | .10 | (1,21) | 10.13 | **.004** | .33 | (1,21) | 28.40 | **< .001** | .58 |
| **Difficulty (easy/hard)** | (1,21) | 41.45 | **< .001** | .66 | (1,21) | 41.80 | **< .001** | .67 | (1,21) | 10.35 | **.004** | .33 |
| **Modality × Difficulty** | (1,21) | 1.05 | .32 | .05 | (1,21) | .01 | .93 | .00 | (1,21) | .50 | .488 | .02 |

**Table S14** Results of repeated-measured ANOVAs for percentage of activation and deactivation voxels thresholded at z=2.3/2.6 in abstract or concrete conditions (Study 3).

| **Threshold z=2.3** | **Activation ANOVA** | | | | **Deactivation ANOVA** | | | |
|---|---|---|---|---|---|---|---|---|
| | *df* | *F* | *p* | $\eta^2$ | *df* | *F* | *p* | $\eta^2$ |
| **concreteness (abstract/concrete)** | (1, 18) | .37 | .55 | .02 | (1, 18) | 13.80 | **.002** | .43 |
| **DMN** | (2, 36) | 17.30 | **< .001** | .49 | (2, 36) | 11.74 | **< .001** | .40 |
| **concreteness × DMN** | (2, 36) | 9.68 | **< .001** | .35 | (2, 36) | 3.37 | .065 | .16 |
| **Threshold z=2.6** | **Activation ANOVA** | | | | **Deactivation ANOVA** | | | |
| | *df* | *F* | *p* | $\eta^2$ | *df* | *F* | *p* | $\eta^2$ |
| **concreteness (abstract/concrete)** | (1, 18) | 3.00 | .100 | .14 | (1, 18) | 9.66 | **.006** | .35 |
| **DMN** | (2, 36) | 14.78 | **< .001** | .45 | (2, 36) | 7.88 | **.001** | .30 |
| **concreteness × DMN** | (2, 36) | 12.88 | **< .001** | .42 | (2, 36) | 2.44 | .118 | .12 |

**Table S15** Results of paired t-tests (Bonferroni corrected for the number of comparisons) between abstract and concrete conditions within each subsystem for activation and deactivation voxels thresholded at z=2.3/2.6 (Study 3).

| **Threshold z=2.3** | **Activation within each subsystem** | | | **Deactivation within each subsystem** | | |
|---|---|---|---|---|---|---|
| | *df* | *t* | *p* | *df* | *t* | *p* |
| **MT** | 18 | -1.05 | .93 | 18 | 2.84 | **.033** |
| **Core** | 18 | -2.15 | .13 | 18 | 3.08 | **.021** |
| **FT** | 18 | 2.57 | **.057** | 18 | 1.85 | .24 |
| **Threshold z=2.6** | **Activation within each subsystem** | | | **Deactivation within each subsystem** | | |
| | *df* | *t* | *p* | *df* | *t* | *p* |
| **MT** | 18 | -.57 | > 1 | 18 | 2.34 | .093 |
| **Core** | 18 | -1.89 | .23 | 18 | 2.34 | .093 |
| **FT** | 18 | 3.34 | **.012** | 18 | 1.50 | .45 |

**Table S16** Results of repeated-measured ANOVAs for percentage of activation and deactivation voxels thresholded at z=2.3/2.6 in emotional and spatial conditions (Study 4).

| **Threshold z=2.3** | **Activation ANOVA** | | | | **Deactivation ANOVA** | | | |
|---|---|---|---|---|---|---|---|---|
| | df | F | p | $\eta^2$ | df | F | p | $\eta^2$ |
| **Cue (emotion/spatial cue)** | (1,25) | 7.35 | **.012** | .23 | (1,25) | .10 | .75 | .004 |
| **DMN** | (2,50) | 14.52 | **< .001** | .37 | (2,50) | 7.15 | **.002** | .22 |
| **Cue × DMN** | (2,50) | 23.37 | **< .001** | .48 | (2,50) | 3.07 | .055 | .11 |
| **Threshold z=2.6** | Activation ANOVA | | | | Deactivation ANOVA | | | |
| | df | F | p | $\eta^2$ | df | F | p | $\eta^2$ |
| **Cue (emotion/spatial cue)** | (1,25) | 7.45 | **.011** | .23 | (1,25) | .24 | .63 | .01 |
| **DMN** | (2,50) | 16.53 | **< .001** | .40 | (2,50) | 6.05 | **.004** | .20 |
| **Cue × DMN** | (2,50) | 22.89 | **< .001** | .48 | (2,50) | 3.07 | .055 | .11 |

**Table S17** Results of paired t-tests (Bonferroni corrected for the number of comparisons) between emotional and spatial conditions within each subsystem for activation and deactivation thresholded at z=2.3/2.6 (Study 4).

| **Threshold z=2.3** | **Activation within each subsystem** | | | **Deactivation within each subsystem** | | |
|---|---|---|---|---|---|---|
| | df | t | p | df | t | p |
| **MT** | 25 | -4.74 | **< .001** | 25 | 1.86 | .23 |
| **Core** | 25 | .20 | >1 | 25 | -.97 | >1 |
| **FT** | 25 | -.34 | >1 | 25 | -.90 | >1 |
| **Threshold z=2.6** | **Activation within each subsystem** | | | **Deactivation within each subsystem** | | |
| | df | t | p | df | t | p |
| **MT** | 25 | -4.51 | **< .001** | 25 | 1.91 | .20 |
| **Core** | 25 | .33 | >1 | 25 | -1.12 | .83 |
| **FT** | 25 | -.26 | >1 | 25 | -.95 | >1 |

**Table S18** Results of repeated-measured ANOVAs for signal change in conflict/no-conflict reading or recall conditions (Study 1).

| | | Signal change | | |
|---|---|---|---|---|
| | | F | p | $\eta^2$ |
| **Main effect** | Task (Reading/Recall) | F(1,28) = 33.92 | **< .001** | .55 |
| | Task Conflict (Pure/Conflict) | F(1,28) = .00 | .99 | .00 |
| | DMN | F(2,56) = 53.59 | **< .001** | .66 |
| **Two-way interaction** | Task × Task Conflict | F(1,28) = 1.44 | .24 | .05 |
| | Task × DMN | F(2,56) = 25.11 | **< .001** | .47 |
| | Task Conflict × DMN | F(2,56) = 7.31 | **.002** | .21 |
| **Three-way interaction** | Task × Task Conflict × DMN | F(2,56) = 4.66 | **.013** | .14 |

**Table S19** Results of repeated-measured ANOVAs for signal change in conflict/no-conflict reading or recall conditions in each DMN subsystem (Study 1).

| | | | **Medial temporal subsystem** | | | **Core subsystem** | | | **Fronto-temporal subsystem** | | |
|---|---|---|---|---|---|---|---|---|---|---|---|
| | | df | F | p | $\eta^2$ | F | p | $\eta^2$ | F | p | $\eta^2$ |
| **Main effect** | Task (Reading/Recall) | 28 | 26.44 | **< .001** | .49 | 78.63 | **< .001** | .74 | 2.79 | .11 | .09 |
| | Task Conflict (Pure/Conflict) | 28 | 2.41 | .13 | .08 | .005 | .94 | .00 | 2.01 | .17 | .07 |
| **Two-way interaction** | Task × Task Conflict | 28 | 5.54 | **.026** | .17 | 1.34 | .26 | .05 | .001 | .97 | .00 |

**Table S20** Results of repeated-measured ANOVAs for signal change in easy/hard verbal or picture conditions (Study 2).

| | | **Signal change** | | |
|---|---|---|---|---|
| | | F | p | $\eta^2$ |
| **Main effect** | Modality (verbal/picture) | F(1,21) = 2.12 | .16 | .09 |
| | Difficulty (easy/hard) | F(1,21) = 1.80 | .19 | .08 |
| | DMN | F(2,42) = 71.43 | **< .001** | .77 |
| **Two-way interaction** | Modality × Difficulty | F(1,21) = 1.32 | .26 | .06 |
| | Modality × DMN | F(2,42) = 30.12 | **< .001** | .59 |
| | Difficulty × DMN | F(2,42) = 12.07 | **< .001** | .37 |
| **Three-way interaction** | Modality × Difficulty × DMN | F(2,42) = .41 | .61 | .02 |

**Table S21** Results of repeated-measured ANOVAs for signal change in easy/hard verbal or picture conditions in each DMN subsystem (Study 2).

| | | | **Medial temporal subsystem** | | | **Core subsystem** | | | **Fronto-temporal subsystem** | | |
|---|---|---|---|---|---|---|---|---|---|---|---|
| | | df | F | p | $\eta^2$ | F | p | $\eta^2$ | F | p | $\eta^2$ |
| **Main effect** | Modality (verbal/picture) | 21 | 9.14 | **.006** | .30 | 12.19 | **.002** | .37 | 21.05 | **< .001** | .50 |
| | Difficulty (easy/hard) | 21 | 4.15 | .054 | .17 | 5.46 | **.029** | .21 | 1.89 | .18 | .08 |
| **Two-way interaction** | Modality × Difficulty | 21 | 1.26 | .28 | .06 | .87 | .36 | .04 | 1.45 | .24 | .07 |

**Table S22** Results of repeated-measured ANOVAs for signal change in relevant-cue/irrelevant-cue abstract or concrete conditions (Study 3).

| | **Signal change** | | | |
|---|---|---|---|---|
| | | F | p | $\eta^2$ |
| **Main effect** | Concreteness (Abstract/concrete) | F(1,18) = 26.19 | **< .001** | .59 |
| | Cue (relevant/irrelevant cue) | F(1,18) = .24 | .63 | .01 |
| | DMN | F(2,36) = 33.55 | **< .001** | .65 |
| **Two-way interaction** | Concreteness × Cue | F(1,18) = 1.43 | .25 | .07 |
| | Concreteness × DMN | F(2,36) = 19.73 | **< .001** | .52 |
| | Cue × DMN | F(2,36) = .63 | .54 | .03 |
| **Three-way interaction** | Concreteness × Cue × DMN | F(2,36) = .97 | .37 | .05 |

**Table S23** Results of repeated-measured ANOVAs for signal change in relevant-cue/irrelevant-cue abstract or concrete conditions in each DMN subsystem (Study 3).

| | | | **Medial temporal subsystem** | | | **Core subsystem** | | | **Fronto-temporal subsystem** | | |
|---|---|---|---|---|---|---|---|---|---|---|---|
| | | df | F | p | $\eta^2$ | F | p | $\eta^2$ | F | p | $\eta^2$ |
| **Main effect** | Concreteness (Abstract/concrete) | 18 | 23.35 | **< .001** | .57 | 35.17 | **< .001** | .66 | 1.54 | .23 | .08 |
| | Cue (relevant/irrelevant cue) | 18 | .06 | .82 | .003 | .008 | .93 | .00 | .92 | .35 | .05 |
| **Two-way interaction** | Concreteness × Cue | 18 | 2.59 | .13 | .13 | .27 | .61 | .02 | .24 | .63 | .01 |

**Table S24** Results of repeated-measured ANOVAs for signal change in no cue (0 cue)/emotional cue (E cue)/spatial cue (S cue)/combined cues (C cues) conditions (Study 4).

| Signal change | | | | |
|---|---|---|---|---|
| | | F | p | $\eta^2$ |
| **Main effect** | Cue (no cue/emotional cue/spatial cue/combined cues) | F(3,75) = 4.11 | **.009** | .14 |
| | DMN | F(2,50) = 45.05 | **< .001** | .64 |
| **Two-way interaction** | Cue × DMN | F(6,150) = 3.45 | **.010** | .12 |

**Table S25** Results of paired t-tests (Bonferroni corrected for the number of comparisons) for signal change between no cue (0 cue)/emotional cue (E cue)/spatial cue (S cue)/combined cues (C cues) in each DMN subsystem (Study 4).

| **Paired t tests** | | **Medial temporal subsystem** | | **Core subsystem** | | **Fronto-temporal subsystem** | |
|---|---|---|---|---|---|---|---|
| | *df* | *t* | *p* | *t* | *p* | *t* | *p* |
| **0 cue vs. E cue** | 25 | .133 | >1 | .53 | .60 | .87 | >1 |
| **0 cue vs. S cue** | 25 | -4.64 | **< .001** | -.36 | .72 | -1.36 | .76 |
| **E cue vs. C cues** | 25 | -4.07 | **< .001** | -.84 | .41 | -2.85 | **.036** |
| **S cue vs. C cues** | 25 | .010 | >1 | .26 | .80 | -.47 | >1 |

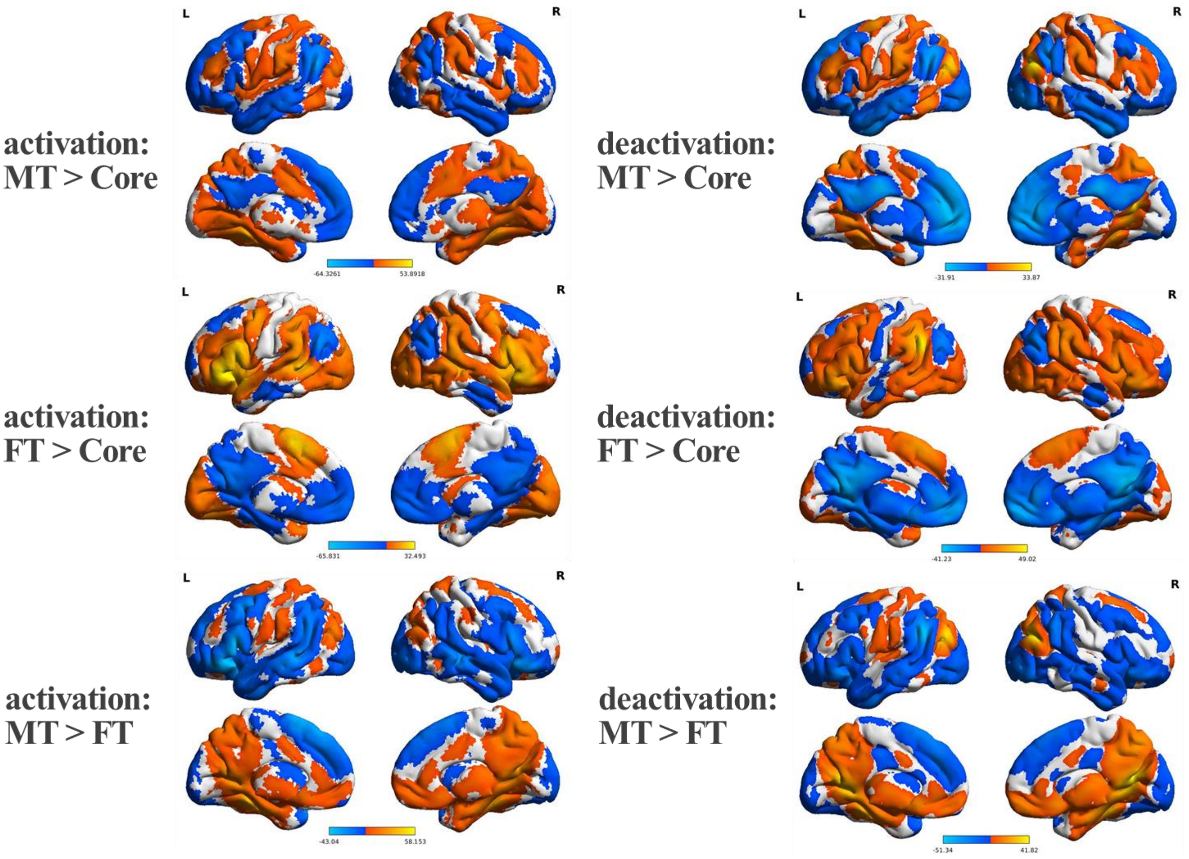


**Figure S5** Contrasts between functional connectivity maps seeding from commonly activated regions in the three subsystems. Each contrast map was FWE-corrected to the voxel-level $p < .001$, and cluster-level $p < .05$. **The upper, middle and lower panels** correspond to the contrasts of MT vs Core, FT vs Core and MT vs FT, respectively. **The left and right panels** correspond to the maps for activation and deactivation seeds respectively.

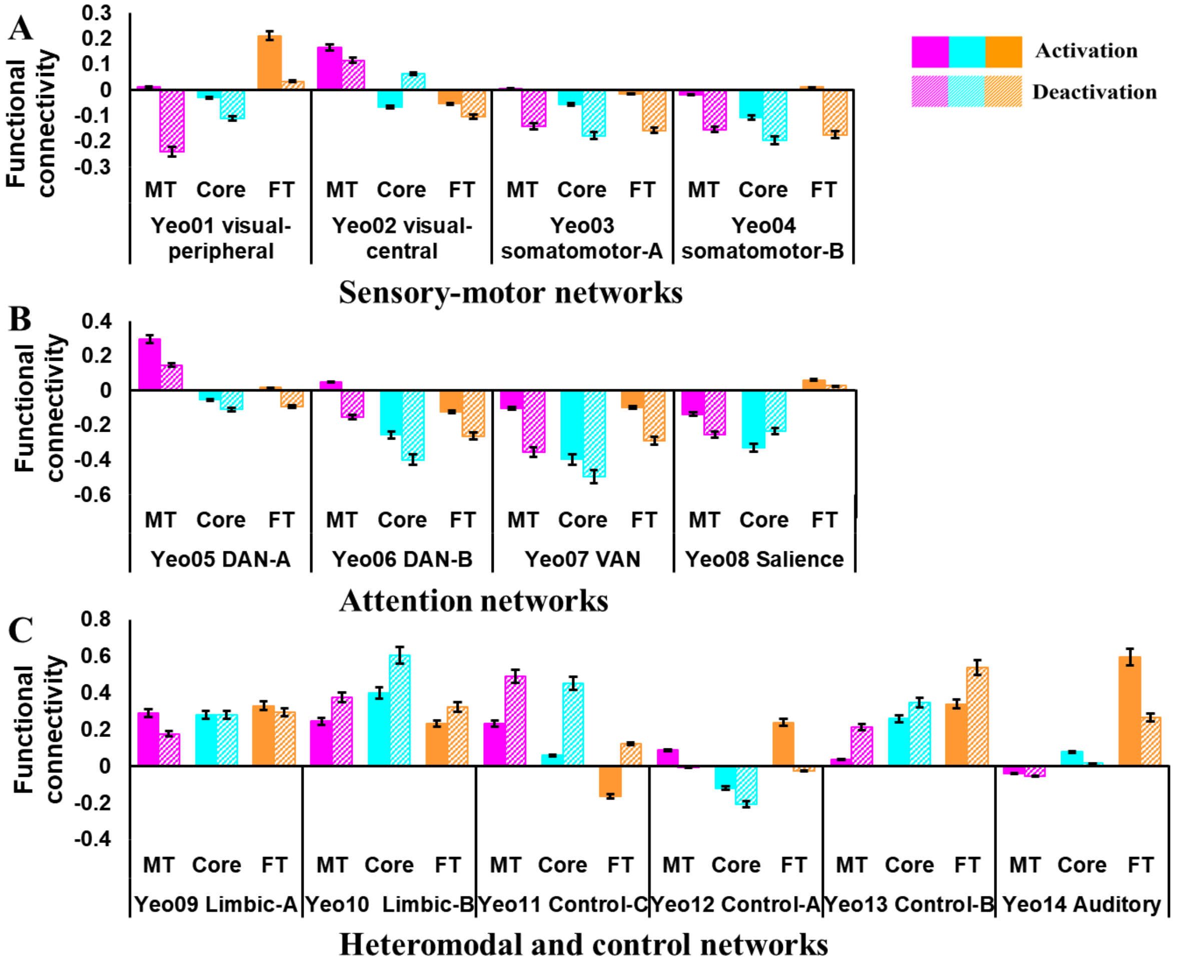


**Figure S6** Functional connectivity between the commonly activated and deactivated regions in DMN subsystems and other large-scale networks (defined by Yeo et al., 2011). **Panels A-C** show the functional connectivity between the commonly activated and deactivated regions in DMN subsystems and (A) sensory-motor networks; (B) attention networks; (C) heteromodal and control networks (pink = medial temporal regions, cyan = core regions, orange = fronto-temporal regions). Solid bars represent the functional connectivity of activation voxels, and grid bars represent the functional connectivity of deactivation voxels. Error bars represent one standard error.
